# Asymmetric distribution of mitochondrial Ca^2+^ regulators specifies compartment-specific mitochondrial function and neuronal development

**DOI:** 10.1101/2025.06.23.660978

**Authors:** Dong Cheol Jang, Su Yeon Kim, Won Seok Kim, Hyunsu Jung, Changuk Chung, Seokyoung Bang, Hong Nam Kim, Hae Woong Choi, Kihoon Han, Yongcheol Cho, Seok-Kyu Kwon

## Abstract

Neuronal polarization is essential for functional compartmentalization, enabling dendritic synaptic integration and axonal action potential generation. While structural differences in mitochondria across compartments have been identified, their functional distinctions remain unclear. Here, we uncovered compartment-specific mitochondrial Ca^2+^ dynamics and their molecular determinants. In axonal mitochondria, Ca^2+^ uptake through MCU occurs independently of ER-stored Ca^2+^ release, with faster matrix Ca^2+^ clearance than dendritic mitochondria, where Ca^2+^ uptake predominantly originates from ER Ca^2+^. The ER-independent mitochondrial Ca^2+^ uptake in axonal mitochondria is mediated by enriched MCU-regulating proteins, MICU1 and MICU2, while higher NCLX expression facilitates rapid Ca^2+^ clearance. Moreover, NCLX knockdown, which functionally mimics a mental retardation-associated mutation, caused more significant axonal branching defects compared to dendrites in vivo, aligning with its enrichment in axons. These findings highlight fundamental Ca^2+^-modulating features and developmental importance of neuronal mitochondria in a compartment-specific manner and reveal the key underlying molecular mechanisms.

## Introduction

Neurons are among the most highly polarized cells in our body, consisting of dendrites, a soma, and a unique axon, crucial for determining the flow of information propagation in the nervous system. The extreme size and morphological polarization of neurons also applies to intracellular organelles such as mitochondria which exhibit drastic structural differences in the axon and dendrite. Axonal mitochondria typically display small (∼1 µm long) and spherical morphologies, whereas dendritic mitochondria adopt elongated, tubular, and highly fused shapes in cortical pyramidal neurons for example ^1, 2^. Moreover, they play pivotal roles in providing energy and regulating Ca^2+^ in both regions, thereby modulating various neuronal functions including synaptic vesicle mobilization and synaptic plasticity ^3–5^. Regarding mitochondrial Ca^2+^ modulation, at presynaptic sites, neurotransmitter release is influenced by mitochondrial Ca^2+^ uptake, with impaired Ca^2+^ import leading to alteration in short-term synaptic plasticity and an increase of asynchronous release ^4, 6–8^. In dendrites of pyramidal neurons, mitochondrial Ca^2+^ regulation also affects multiple synaptic features including excitability and spine dynamics. Despite these compartment-specific roles, whether mitochondrial Ca^2+^ regulation exhibits distinct features in axons and dendrites remains largely unexplored.

Recent studies have elucidated multiple components involved in mitochondrial Ca^2+^ influx and efflux. The mitochondrial Ca^2+^ uniporter (MCU), a channel responsible for rapid Ca^2+^ uptake, forms a complex with other subunits such as mitochondrial Ca^2+^ uptake 1 (MICU1), MICU2, or MICU3, acting as gatekeepers and modifiers of the Ca^2+^ uptake properties of MCU ^9–11^. Conversely, mitochondrial Ca^2+^ clearance from the mitochondrial matrix is regulated by the Na^+^/Ca^2+^ exchanger (NCLX) under physiological conditions. Notably, the composition of these components can vary across tissues. For example, the MICU1:MCU expression ratio is lower in heart and skeletal muscle compared to the liver, leading to lesser threshold and cooperativity of MCU activation ^12^. Additionally, in the brain, mitochondrial Ca^2+^ regulators show distinct expression patterns depending on subregions; for instance, there is a higher level of MCU in hippocampal CA2/3 than CA1 ^13, 14^. However, whether these regulatory components are differentially distributed or tuned across axonal and dendritic compartments remains unknown.

Mitochondrial dysfunction has long been implicated in neurodegenerative diseases, where defects in energy metabolism and Ca^2+^ buffering contribute to progressive neuronal loss ^11, 15–17^. In addition, several studies have also shown that mitochondrial defects can interfere with key processes in neuronal development, including axon branching and dendritic arborization ^18, 19^. Given that mitochondria serve distinct roles in axons and dendrites, their dysregulation could differentially affect these subcellular compartments depending on local functional demands. However, whether mitochondrial dysfunction divergently impacts axonal and dendritic development remains unclear.

In this study, we investigated the functional asymmetry of mitochondria in axons and dendrites regarding Ca^2+^ regulation and identified key molecular components responsible for these differences following axon isolation. Furthermore, we explored the importance of the compartment-specific functional properties of mitochondria regarding on neuronal development.

## Results

### Axonal mitochondria exhibit faster Ca^2+^ decay compared to dendritic mitochondria in cortical pyramidal neurons

Recent advances in Ca^2+^ monitoring techniques have enabled the direct measurement of mitochondrial Ca^2+^ dynamics using mitochondria-targeted genetically encoded Ca^2+^ sensors ^20^. We generated mitochondria-targeted-jGCaMP8m (mito-jGCaMP8m) by incorporating the mitochondria targeting sequence of COXVIII and transfected it with mScarlet in cortical pyramidal neurons using *ex utero* electroporation (EUE) at E15.5 (Extended Data Fig. 1a). The *ex utero* electroporation method selectively labels cortical pyramidal neurons by introducing plasmids to neural progenitors lining the lateral ventricles ^21^. Following dissociated culture for 15-21 days *in vitro* (DIV), mitochondrial Ca^2+^ dynamics in dendrites and axons were monitored following 10 action potentials (APs) stimulation. In order to establish stable baselines during recording, while leaving evoked activity intact, we added low concentrations of tetrodotoxin (TTX, less than 50nM) to block spontaneous activity only (Extended Data Fig. 2). To ensure comparability of mitochondrial Ca^2+^ dynamics within each compartment at similar Ca^2+^ levels, we locally applied electrical field potentials using a glass pipette containing an electrode near distal dendrites or axons (Extended Data Fig. 1b).

We first assessed mitochondrial Ca^2+^ release properties between axons and dendrites post-stimulation (10 AP at 10 Hz). Surprisingly, following rapid Ca^2+^ uptake, axonal mitochondria exhibited a decay time constant twice as fast as that of dendritic mitochondria (20 sec for axons, 40 sec for dendrites. Fig. 1a-d and Supplementary Video 1). We did not observe any significant correlation between the peak of mitochondrial matrix Ca^2+^ amplitude and the decay time under these conditions (Extended Data Fig. 1c). These findings suggest an intrinsic disparity in mitochondrial Ca^2+^ extrusion properties between axons and dendrites.

**Fig. 1.**
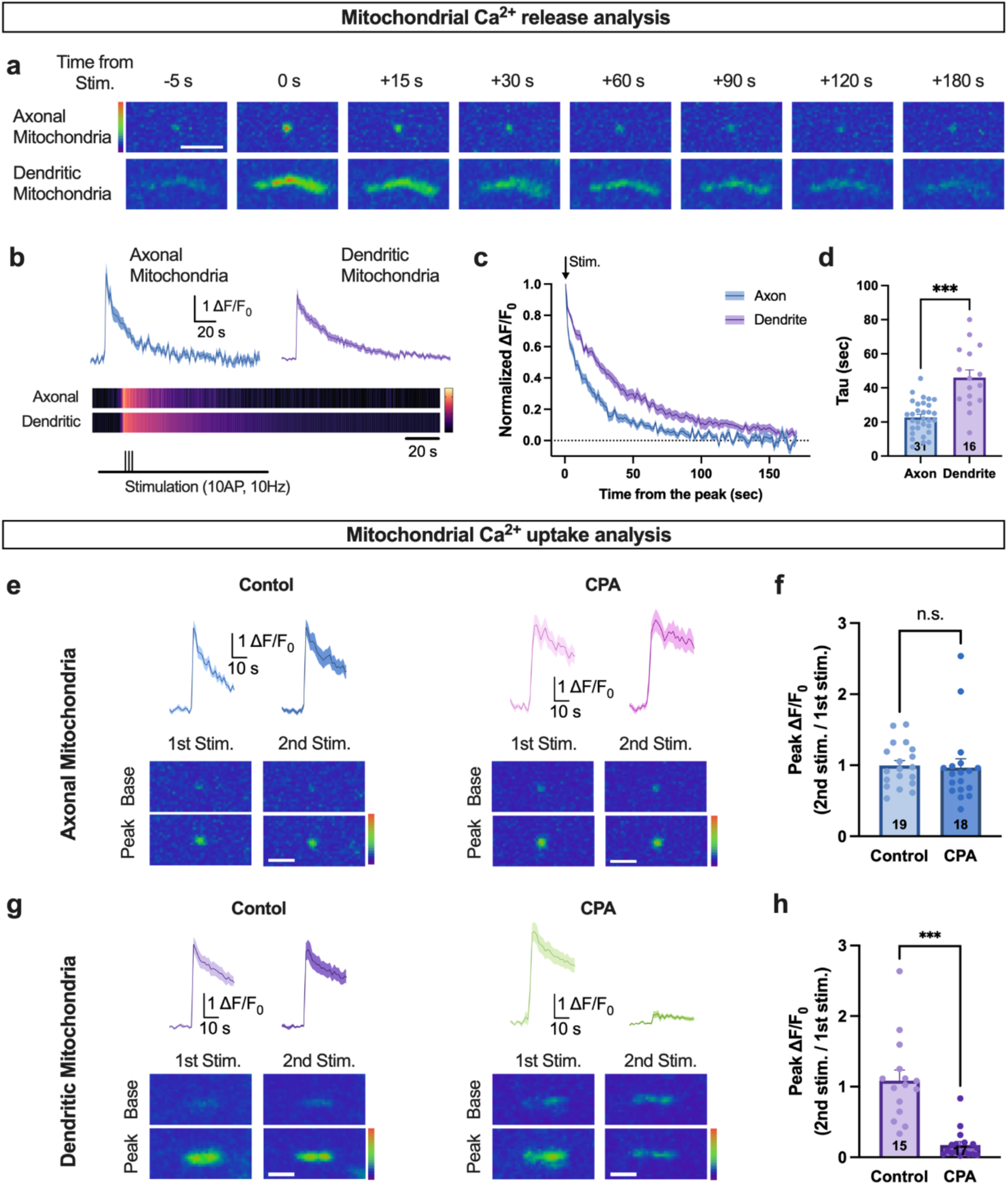
Differential Ca^2+^ decay and uptake between axonal and dendritic mitochondria in cortical pyramidal neurons. (a) Time series images from axonal and dendritic mitochondria. (b) Representative Ca^2+^ traces and line plots. (c) Normalized ΔF/F_0_ traces by peak value. (d) Calculated decay tau (τ) values. Axonal mitochondria exhibited significantly faster Ca^2+^ decay than dendritic mitochondria (Axon n=31 mitochondria from 31 axons, 22.7±1.72, Dendrite n=16 mitochondria from 16 dendrites, 46.0±4.50, p<0.001). (e) Ca^2+^ responses after the first and the second stimulation from axonal mitochondria (see also Extended Data Fig. 3a). Representative Ca^2+^ traces and images from the control group (left) and CPA-treated group (right). (f) Both groups showed similar Ca^2+^ responses after the first and the second stimulation. No differences were observed in normalized peak ΔF/F_0_ values by the first peak value between the control and CPA-treated group (Control n=19 mitochondria from 19 axons, 0.996±0.0702, CPA n=18 mitochondria from 18 axons, 0.967±0.124, p=0.835). (g) Ca^2+^ responses after the first and the second stimulation from dendritic mitochondria. Representative Ca^2+^ traces and images from the control group (left) and CPA-treated group (right). (h) Control groups showed similar Ca^2+^ responses after the first and the second stimulation, whereas the CPA-treated group showed a significantly reduced Ca^2+^ response after the second stimulation. Ca^2+^ responses at the second stimulation in both groups were significantly different (Control n=15 mitochondria from 15 dendrites, 1.09±0.189, CPA n=17 mitochondria from 17 dendrites, 0.231±0.0635, p<0.001). Axonal and dendritic mitochondrial Ca^2+^ dynamics were monitored using mito-jGCaMP8m with mScarlet at 15-21 DIV. Unpaired t-test was used for bar graphs. Mitochondrial numbers are annotated at the bottom of bar graphs. Data are presented as mean ± SEM. ***p < 0.001. n.s., not significant. Scale bar, 5µm.

### Dendritic, but not axonal, mitochondrial Ca^2+^ uptake is largely dependent on ER-stored Ca^2+^ release in cortical pyramidal neurons

In dendrites, mitochondrial Ca^2+^ uptake relies heavily on Ca^2+^ released from the ER following synaptic stimulation ^22^. However, in hippocampal neurons, unlike dendritic ER, axonal ER is capable of importing Ca^2+^ upon action potentials ^23^. To compare mitochondrial Ca^2+^ uptake properties between axons and dendrites while minimizing the influence of ER, we added cyclopiazonic acid (CPA), which inhibits the sarco/endoplasmic reticulum Ca^2+^-ATPase (SERCA) pump, thus depleting ER-stored Ca^2+^. Moreover, to ensure clear depletion of ER-stored Ca^2+^ in the absence of spontaneous activity by treating a small amount of TTX, we applied 2 stimulus trains (10 APs at 10 Hz) within 7 min interval (Extended Data Fig. 3a). Stimulation of distal dendrites was not enough for inducing considerable amount of ER Ca^2+^ release, therefore the electrode was placed nearby cell bodies (Extended Data Fig. 3b).

Axonal mitochondria of cortical pyramidal neurons remain unaffected by the addition of CPA (Fig. 1e, f and Supplementary Video 2), but on the contrary, dendritic mitochondrial Ca^2+^ uptake is significantly reduced (∼77%) under CPA-treated conditions in agreement with a previous report ^22^, suggesting that dendritic mitochondria inherently possess lower Ca^2+^ uptake capabilities (Fig. 1g,h and Supplementary Video 3). This is not caused by difference in cytosolic Ca^2+^ levels, although dendritic ER-stored Ca^2+^ release is successfully depleted following CPA treatment (Extended Data Fig. 3c-e). This could be because most ER-released Ca^2+^ might be transferred to the mitochondria, or cytosolic Ca^2+^ from the ER upon stimulation could reduce the Ca^2+^ gradient across the plasma membrane, thereby affecting extracellular Ca^2+^ import. Overall, these results indicate the mitochondrial Ca^2+^ uptake properties are also different in dendrites and axons; axonal mitochondrial Ca^2+^ uptake capacity upon action potentials is significantly higher than that of dendritic mitochondria.

### Mitochondrial Ca^2+^ regulatory components exhibit differential expression between dendrites and axons of cortical neurons

To explore the molecular determinants underlying the functional differences in Ca^2+^ dynamics between axonal and dendritic mitochondria, cortical neurons were cultured on a 3 μm porous membrane, allowing isolation of axons penetrating beneath the membrane (Fig. 2a). In addition, these cultures were incubated in the media without FBS to minimize the effect of astrocytes (Extended Data Fig. 4a). The whole neuron fraction above the membrane was collected using a cell scraper, followed by submersion of the membrane in the lysis buffer to extract the remaining axonal components. To validate the relative enrichment of axonal fraction to the whole neuron fraction, the levels of Histone H3 for the cell body, synaptophysin for the axon, and GluR2 for the dendrite were examined. As anticipated, Histone H3 and GluR2 levels were significantly lower in the axon fraction, whereas the presynaptic vesicle protein was enriched (Fig. 2b-e).

**Fig. 2.**
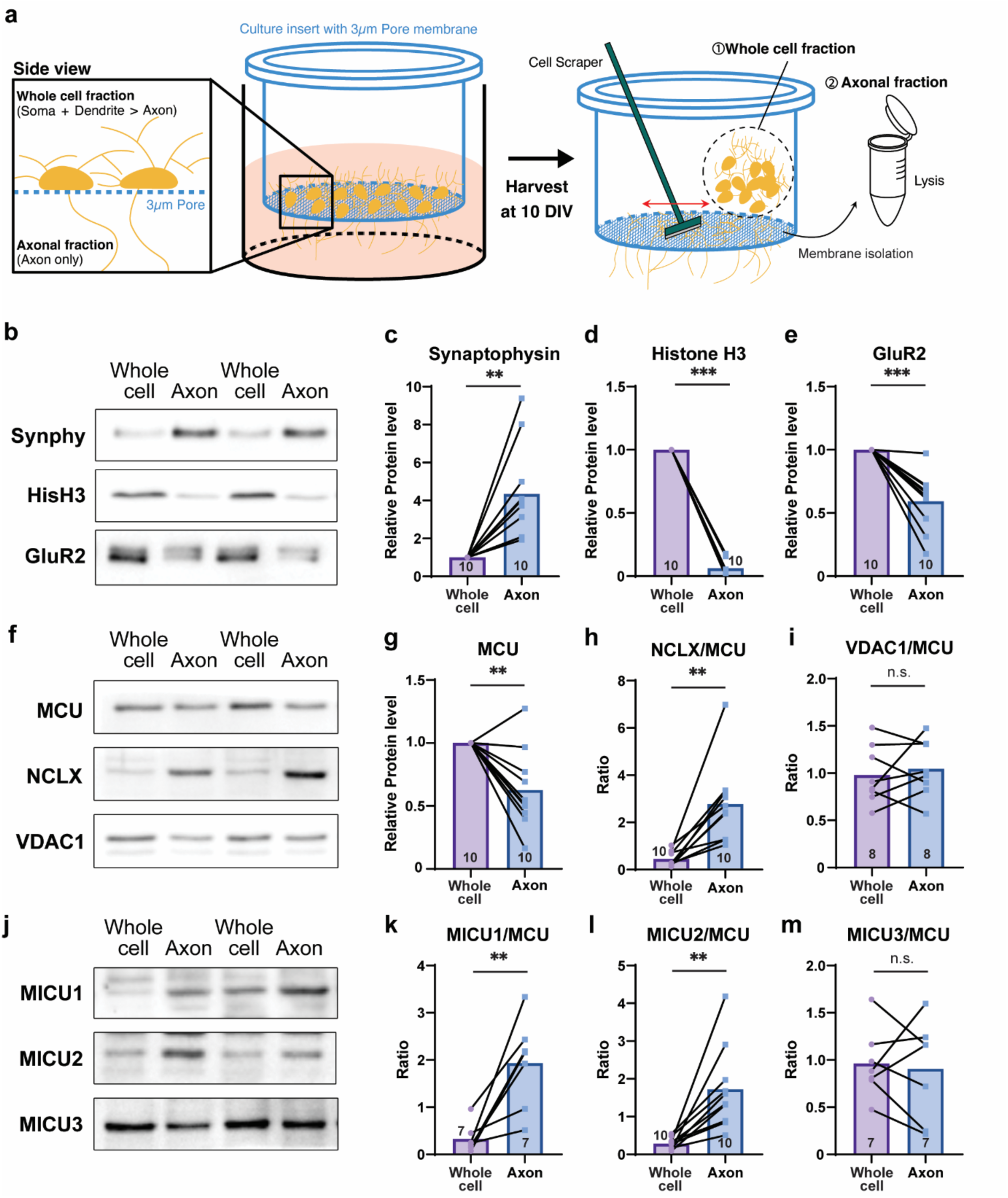
NCLX, MICU1, and MICU2 proteins are enriched in axonal fraction. (a) Schematic representations of harvesting whole cell fractions and axonal fractions separately. (b-e) Separation of whole cell and axonal fraction is confirmed by synaptophysin (Synphy, presynapse/axon), histone H3 (HisH3, nuclear/cell body), and GluR2 (postsynapse/dendrite) levels. Synaptophysin level is significantly higher in the axonal fraction, but not histone H3 and GluR2. Representative Western blot images (b) and bar graph showing relative protein level of synaptophysin (c, Axon 4.350±0.799, p=0.002), Histone H3 (d, Axon 0.063±0.024, p<0.001) and GluR2 (e, Axon 0.594±0.0713, p<0.001). (f-j) Western blots of mitochondrial Ca^2+^ regulatory components in the whole cell and the axonal fraction. These protein levels were normalized by MCU. Significantly higher ratios of NCLX, MICU1, and MICU2 to MCU are shown in the axon, while the ratios of VDAC1/MCU and MICU3/MCU are similar. Representative western blot image (f, j) and bar graph showing relative protein level of MCU (g, Axon 0.627±0.100, p=0.005) and X:MCU ratio of NCLX (h, Whole cell 0.452±0.102, Axon 2.779±0.540, p<0.001), VDAC1 (i, Whole cell 0.979±0.109, Axon 1.047±0.106, p=0.551), MICU1 (k, Whole cell 0.329±0.115, Axon 1.932±0.355, p=0.005), MICU2 (l, Whole cell 0.284±0.048, Axon 1.724±0.346, p=0.002) and MICU3 (m, Whole cell 0.963±0.138, Axon 0.905±0.199, p=0.751). Relative protein levels were normalized to values from the whole cell fraction. Paired t-test was used for bar graphs. Sample numbers are annotated at the bottom of bar graphs. Data are presented as mean ± SEM. **p < 0.01. ***p < 0.001. n.s., not significant.

Subsequently, we investigated whether mitochondrial Ca^2+^ regulatory proteins differ between axonal and somatodendritic compartments. Given that axonal mitochondria are smaller in size and less dense than in the somatodendritic region ^1, 2^, the total mitochondrial protein content of the axon may be lower than that of dendrites when the same protein quantity is loaded. To test this, cortical pyramidal neurons expressing mito-YFP and HA-mCherry were immunostained with MCU antibody and intensities of MCU were similar in the dendritic and axonal mitochondria at 10 DIV (Extended Data Fig. 4h), even though the Western blot intensity is higher in whole neuron fraction (Fig. 2f,g). Antibodies for other Ca^2+^ regulatory proteins are not available for immunocytochemistry; therefore, to mitigate this potential side effect, the protein X:MCU ratio was assessed.

Surprisingly, the mitochondrial Ca^2+^ release protein NCLX shows a significantly higher expression in the axon, consistent with the faster decay times of axonal mitochondria (Fig 2h, Extended Data Fig. 4g). Regarding mitochondrial Ca^2+^ uptake, the ratio of MICUs to MCU is critical for the uptake capacity, with higher MICU1/2 to MCU increasing the opening threshold while cooperatively importing more Ca^2+^ than lower MICU1/2-associated MCU complexes ^12, 24–26^. VDAC1 is a non-selective channel passing ions, nucleotides, and metabolites in the outer mitochondrial membrane (OMM), and MICU3 is known to be enriched in neurons enhancing the MCU activity ^23, 27^. Interestingly, significantly higher ratios of MICU1/MCU and MICU2/MCU were observed in the axonal fraction, although VDAC1 and MICU3 to MCU ratios were similar between the whole cell and the axon fraction (Fig. 2i-m). Non-normalized TOM40, VDAC1, and MICU3 levels were lower in the axon, like the MCU immunoblot, due to a lower mitochondrial amount as mentioned above, even though NCLX, MICU1, and MICU2 amounts were still more in the axon (Extended Data Fig. 4b-g). These results clearly show that NCLX, involved in mitochondrial Ca^2+^ release, and MICU1 and MICU2, related to mitochondrial Ca^2+^ uptake, are significantly enriched in axonal mitochondria.

### Neuronal compartment-specific mitochondrial Ca^2+^ dynamics and related protein expression profiles are conserved in hippocampal neurons

Since the initial findings were based solely on cortical neurons, we next investigated whether the compartment-specific differences in mitochondrial Ca²⁺ regulation represent a generalizable feature of neuronal architecture or a neuron type-dependent characteristic. To address this, we examined mitochondrial Ca^2+^ dynamics of hippocampal pyramidal neurons following *ex utero* hippocampal electroporation with mito-jGCaMP8m and mScarlet. Similar with cortical neurons, axonal mitochondria exhibited a significantly faster Ca²⁺ decay than dendritic mitochondria, with average decay time constants of approximately 60 and 130 seconds, respectively (Fig. 3a–c). Moreover, Western blot analysis of isolated axons and whole cell fractions revealed that mitochondrial Ca²⁺ extrusion protein NCLX, along with the uptake modulators MICU1 and MICU2, were enriched in the axons of hippocampal neurons, whereas proteins such as MICU3 was expressed at comparable levels across compartments and VDAC1 showed slightly lower amount in axons (Fig. 3d–i, Extended Data Fig. 5). Taken together, these findings indicate that the neuronal compartment-specific differences in mitochondrial Ca²⁺ dynamics and associated protein expression are largely conserved features across distinct brain regions.

**Fig. 3.**
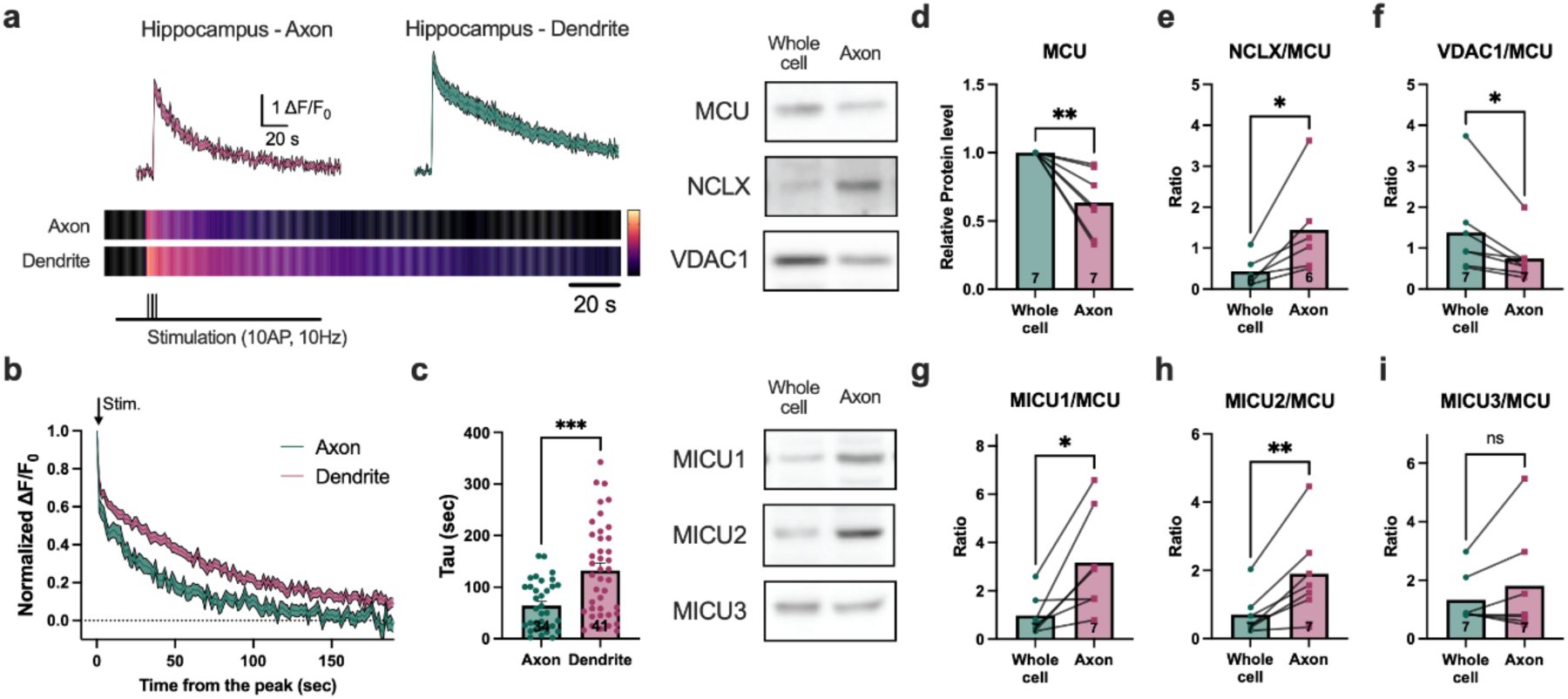
Mitochondrial Ca^2+^ dynamics and associated-protein expression patterns are mostly conserved in hippocampal neurons. Axonal mitochondria exhibited significantly faster Ca^2+^ decay than dendritic mitochondria. (a) Representative Ca^2+^ traces and corresponding line plots. (b) ΔF/F_0_ traces normalized to peak value. (c) Calculated decay tau (τ) values. (d-i) Western blots of mitochondrial Ca^2+^ regulatory components in the whole cell and the axonal fraction. Protein levels were normalized with MCU. NCLX, MICU1, and MICU2 showed significantly higher expression relative to MCU in the axonal fraction, while MICU3/MCU ratio is similar and VDAC1/MCU is slightly lower. Bar graph showing relative protein level of MCU (d, Axon 0.634±0.090, p=0.006) and X:MCU ratio of NCLX (e, Whole cell 0.434±0.148, Axon 1.443±0.471, p=0.036), VDAC1 (f, Whole cell 1.381±0.421, Axon 0.749±0.218, p=0.027), MICU1 (g, Whole cell 0.970±0.316, Axon 3.172±0.815, p=0.017), MICU2 (h, Whole cell 0.704±0.238, Axon 1.897±0.497, p=0.006) and MICU3 (i, Whole cell 1.330±0.327, Axon 1.809±0.692, p=0.253). Relative protein levels were normalized to values from the whole cell fraction. Paired t-test. Sample numbers are annotated at the bottom of bar graphs. Data are presented as mean ± SEM. **p < 0.01. ***p < 0.001. n.s., not significant.

### The differential mitochondrial Ca^2+^ uptake capacity between dendrites and axons is determined by axon-enriched MICU1 and MICU2

Then, we checked if the lower amount of MICU1 and MICU2 in dendrites results in reduced Ca^2+^ uptake ability compared to axonal mitochondria. To address this, we overexpressed MICU1 or MICU2 in cortical pyramidal neurons using *ex utero* electroporation, then measured mitochondrial Ca^2+^ dynamics with CPA treatment (Fig. 4, Extended Data Fig. 6a). Axonal mitochondria displayed no significant changes in the normalized peak level (2^nd^ stimulation/1^st^ stimulation) upon MICU1 or MICU2 overexpression (Fig. 4a, b and Supplementary Video 4), although absolute peak value is significantly lower in MICU1 overexpression, but slightly higher in MICU2 overexpression (Fig. 4a and Extended Data Fig. 6b). Decreased mitochondrial Ca^2+^ level by MICU1 overexpression may be caused from tightening of cristae junction, which could lead reduction of cristae Ca^2+^ influx ^28^. In contrast, dendritic mitochondrial Ca^2+^ uptake was dramatically increased upon the same stimulation condition (Fig. 4c, d, Extended Data Fig. 6c and Supplementary Video 5), suggesting that MICU1 or MICU2 is sufficient to cooperatively enhance MCU-mediated Ca^2+^ import. Since MICU2 interacts with MCU via MICU1^25, 26, 29, 30^, MICU2 overexpression alone has a weaker effect on altering Ca^2+^ regulatory function. Taken together, axonally-enriched MICU1/2 not only enhance mitochondrial Ca^2+^ uptake but also grant ER-independency, suggesting that MICU1/2 serve as key determinants for mitochondrial Ca^2+^ uptake modality.

**Fig. 4.**
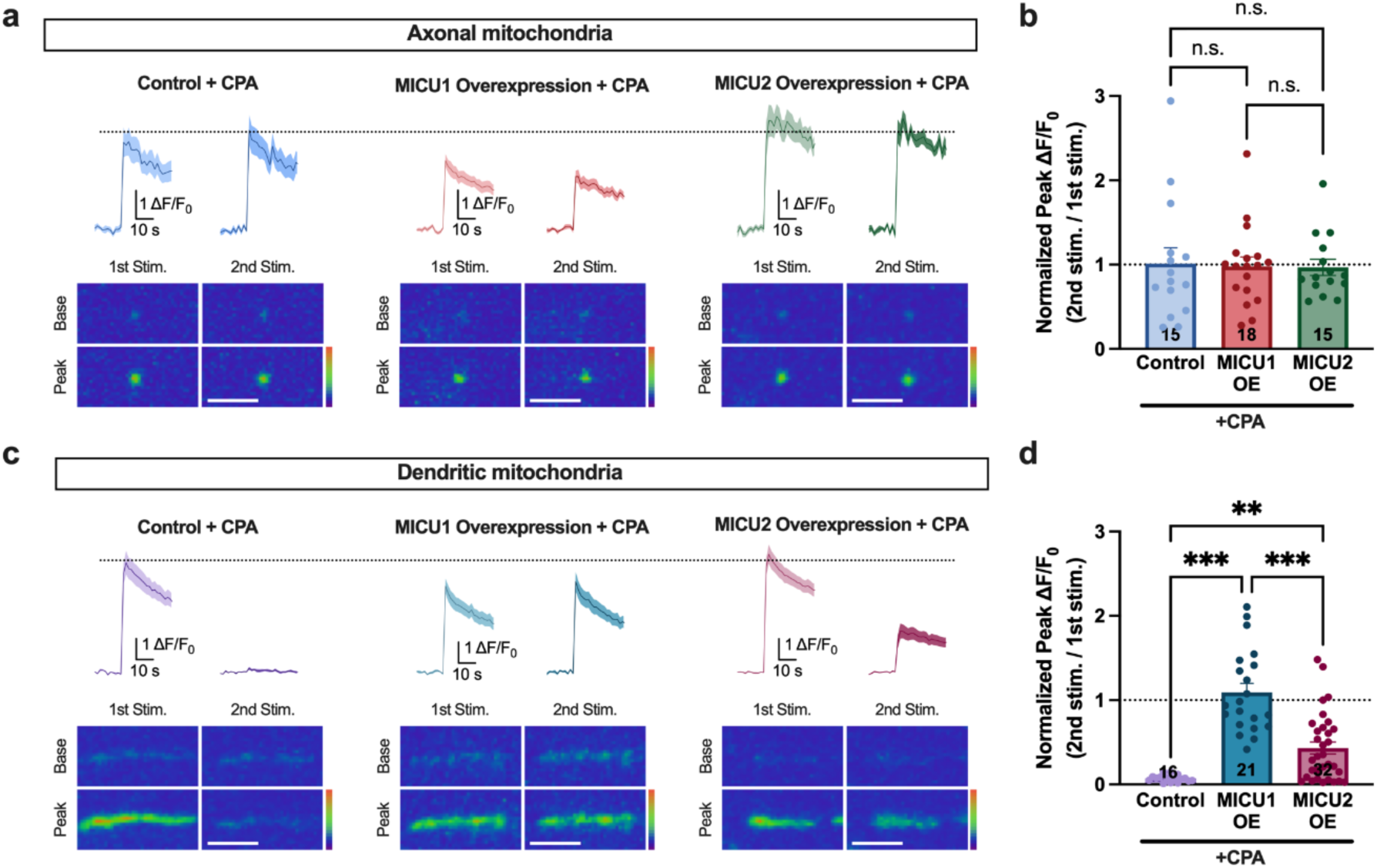
MICU1 or MICU2 overexpression is sufficient for ER-independent Ca^2+^ import of dendritic mitochondria. (a) Overexpression of MICU1 or MICU2 did not affect the Ca^2+^ uptake of axonal mitochondria. Representative Ca^2+^ traces and images of axonal mitochondria from the control group (left), MICU1 overexpression group (middle), and MICU2 overexpression group (right). (b) No differences were found in normalized peak ΔF/F_0_ values (peak from 2^nd^ stimulation/peak from 1^st^stimulation) between the three groups (One-way ANOVA, F=0.0276, p=0.973; Control n=15 mitochondria from 15 axons, 1.01±0.187, MICU1 OE n=18 mitochondria from 18 axons, 0.979±0.112, MICU2 OE n=15 mitochondria from 15 axons, 0.967±0.0967, Control vs MICU1 OE p=0.984, Control vs MICU2 OE p=0.972, MICU1 OE vs MICU2 OE p=0.998). (c) Overexpression of MICU1 increased the Ca^2+^ uptake capacity of dendritic mitochondria and made it fully independent from ER-released Ca^2+^, while overexpression of MICU2 showed partial enhancement. Representative Ca^2+^ traces and images of dendritic mitochondria from the control group (left), MICU1 overexpression group (middle), and MICU2 overexpression group (right). (d) Normalized peak graph between the three groups. Both MICU1 and MICU2 overexpression groups showed a significant increase in normalized peak value. But the increases in the MICU1 overexpression group were significantly larger than the MICU2 overexpression group (One-way ANOVA, F=34.3, p<0.001; Control n=16 mitochondria from 16 dendrites, 0.0704±0.0102, MICU1 OE n=21 mitochondria from 21 dendrites, 1.09±0.107, MICU2 OE n=32 mitochondria from 32 dendrites, 0.432±0.0710; Control vs MICU1 OE p<0.001, Control vs MICU2 OE p=0.009, MICU1 OE vs MICU2 OE p<0.001). Axonal and dendritic mitochondrial Ca^2+^ dynamics of control, MICU1 overexpressing, or MICU2 overexpressing cortical pyramidal neurons were monitored using mito-jGCaMP8m at 15-21 DIV. All recordings were performed under 30 µM CPA treatment. One-way ANOVA with *post hoc* Tukey’s test was used for panels B and D. Mitochondrial numbers are annotated at the bottom of bar graphs. Data are presented as mean ± SEM. **p < 0.01. ***p < 0.001. n.s., not significant. Scale bar, 5µm.

### MICU1 overexpression increases the vulnerability of dendritic mitochondria to excitotoxic stress

Previous study showed that increased mitochondrial Ca^2+^ uptake by MCU overexpression exacerbates excitotoxic cell death due to rapid loss of mitochondrial membrane potential following NMDA treatment ^31^. ER-stored Ca^2+^ serves as the primary Ca^2+^ source for dendritic mitochondria, with resting ER Ca^2+^ levels reaching up to 1 mM ^32, 33^. Therefore, a massive influx of Ca^2+^ released from the ER and imported from the extracellular space, combined with elevated mitochondrial Ca^2+^ uptake due to MICU1 overexpression, could lead to rapid mitochondrial Ca^2+^ overload, potentially damaging them.

To investigate this, we examined changes in dendritic mitochondrial membrane potential under excitotoxic conditions in neurons overexpressing MICU1. Mitochondrial membrane potential was monitored using tetramethylrhodamine methyl ester (TMRM) in response to NMDA treatment (20 µM) in both control and MICU1-overexpressing neurons. Membrane potential, represented as the F_mitochondria_/F_cytoplasm_ (F_m_/F_c_), declined more rapidly in the MICU1 overexpression group than the control group (Extended data Fig. 7a). F_m_/F_c_ values at 1, 5, and 10 minutes post-stimulation were significantly lower in MICU1-overexpressing dendritic mitochondria, with no difference observed before treatment (Extended data Fig. 7b,c). These findings indicate that MICU1 overexpression induces Ca^2+^ overload in dendritic mitochondria following NMDA treatment, increasing susceptibility to excitotoxicity. Based on these results, we conclude that low MICU1 expression in dendritic mitochondria limits the Ca^2+^ source to the ER, thus protecting mitochondria from excitotoxic stress.

### Faster axonal mitochondrial Ca^2+^ release is regulated by higher levels of NCLX compared to dendritic mitochondria

To investigate whether the elevated amount of NCLX in the axon contributes to its faster Ca^2+^ release property, we introduced a NCLX knockdown plasmid co-expressing mTagBFP2, along with mito-jGCaMP8m and mScarlet as a cell filler (Extended Data Fig. 8). We then compared the axonal mitochondrial Ca^2+^ decay constant with that of dendrites following 10AP stimulation. Under NCLX-deficient condition, axonal mitochondria exhibited an extended Ca^2+^ release time, reaching a level similar to that of dendritic mitochondria (Fig. 5b, c and Supplementary Video 6, 7). Interestingly, dendritic mitochondria showed no significant changes in the partial knockdown condition (Fig. 5b, c and Extended Data Fig. 8a); however, it appears to have a more pronounced impact on axonal mitochondrial function (Fig. 5). These results suggest that the rapid extrusion of axonal mitochondrial Ca^2+^ is derived from the higher levels of NCLX.

**Fig. 5.**
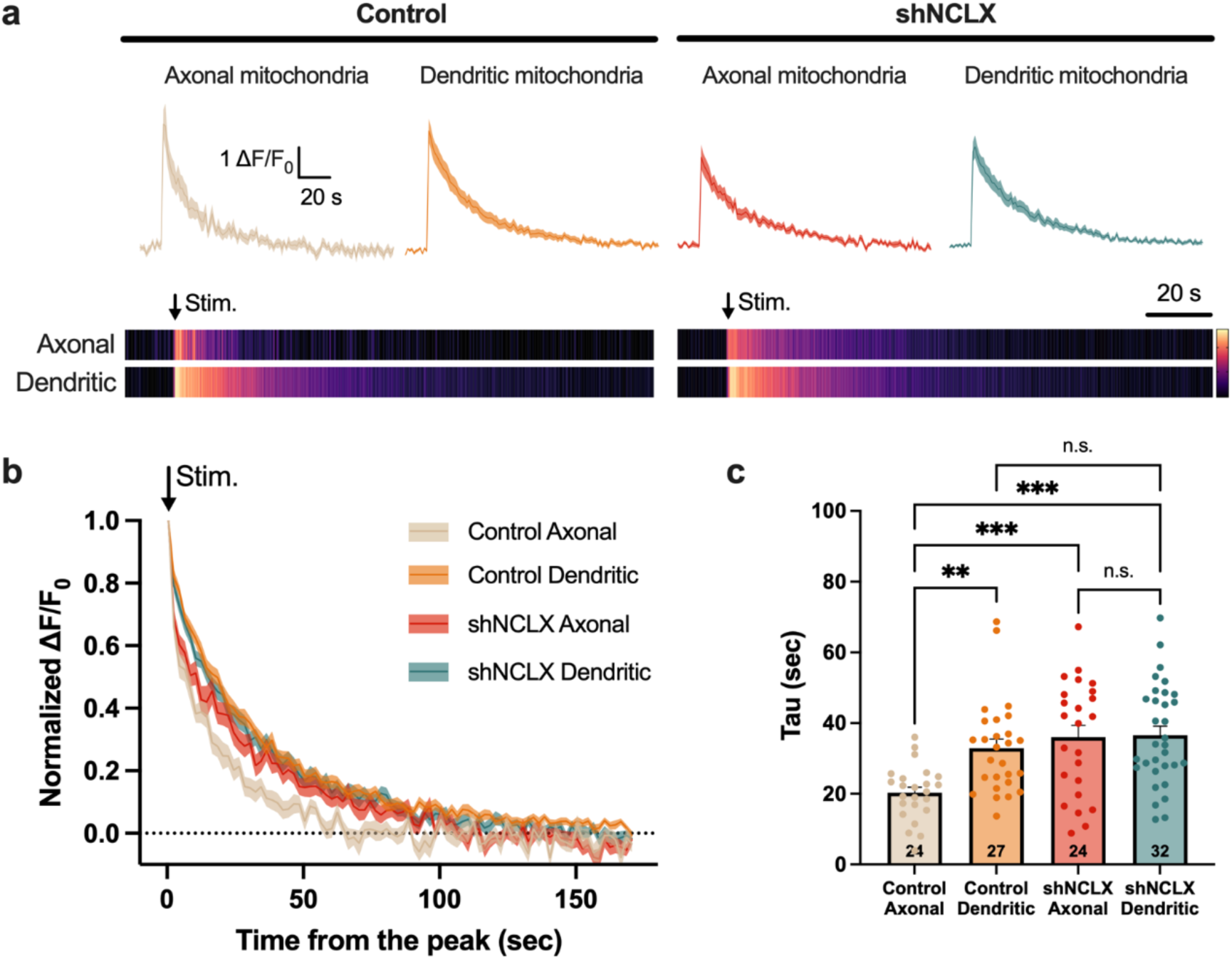
Faster Ca^2+^ efflux in axonal mitochondria attributed to elevated NCLX levels relative to dendritic mitochondria. (a) Representative Ca^2+^ traces of the control group (left) and shNCLX (NCLX knockdown) group (right). (b) Normalized ΔF/F_0_ traces by peak value. NCLX knockdown affects only the Ca^2+^ release of axonal mitochondria. (c) Calculated decay tau (τ) values. NCLX knockdown significantly increases decay tau of axonal mitochondria, but not in dendritic mitochondria (One-way ANOVA, F=8.25, p<0.001; Control Axon n=24 mitochondria from 24 axons, 20.3±1.57, Control Dendrite n=27 mitochondria from 27 dendrites, 32.9±2.53, shNCLX Axon n=24 mitochondria from 24 axons, 36.0±3.31, shNCLX Dendrite n=32 mitochondria from 32 dendrites, 36.6±2.52, Control Axon vs Control Dendrite p=0.005, Control Axon vs shNCLX Axon p<0.001, Control Axon vs shNCLX Dendrite p<0.001, Control Dendrite vs shNCLX Axon p=0.839, Control Dendrite vs shNCLX Dendrite p=0.717, shNCLX Axon vs shNCLX Dendrite p=0.999). Axonal and dendritic mitochondrial Ca^2+^ dynamics of control or NCLX knockdown cortical pyramidal neurons were monitored using mito-jGCaMP8m at 15-21 DIV. One-way ANOVA with *post hoc* Tukey’s test was used for panel c. Mitochondrial numbers are annotated at the bottom of a bar graph. Data are presented as mean ± SEM. **p < 0.01. ***p < 0.001. n.s., not significant.

### NCLX knockdown significantly impairs axon branching, but not dendritic arborization

Mutation or reduced expression of NCLX have been linked to mental retardation and neurodegenerative conditions including Alzheimer’s disease ^34, 35^. Especially, NCLX P367S mutation near the catalytic domain, which is identified from severe mental retarded patients, shows similar mitochondrial Ca^2+^ dynamics with NCLX KO condition ^35^. Therefore, we tested whether NCLX-deficiency can affect neuronal development in a compartment-dependent manner. To this end, we performed unilateral *in utero* electroporation at E15.5 to introduce control (scrambled shRNA) or shNCLX together with an mScarlet as a neuronal filler into layer 2/3 cortical pyramidal neurons and assessed their morphologies at P21. In control brains, callosal axons extended across the midline and formed a dense and stereotyped terminal branching pattern in the contralateral side, targeting both layer 2/3 and layer 5 (Fig. 6a–c). In contrast, NCLX knockdown led to a marked and layer-specific reduction in axonal branching, particularly in layer 2/3, despite normal trajectory and targeting (Fig. 6d–g). To check the impact on dendrites, we reconstructed the dendritic morphologies from the ipsilateral part of the electroporated cortex. A mild reduction in complexity was noted in the shNCLX group, but the difference was not statistically significant (Fig. 6h–l), indicating that dendritic arborization is relatively resilient to NCLX deficiency. Together, these findings demonstrate that the asymmetric expression of NCLX leads to differential compartmental requirements during neuronal development, with axons being more sensitive to NCLX-dependent mitochondrial Ca^2+^ regulation than dendrites.

**Fig 6.**
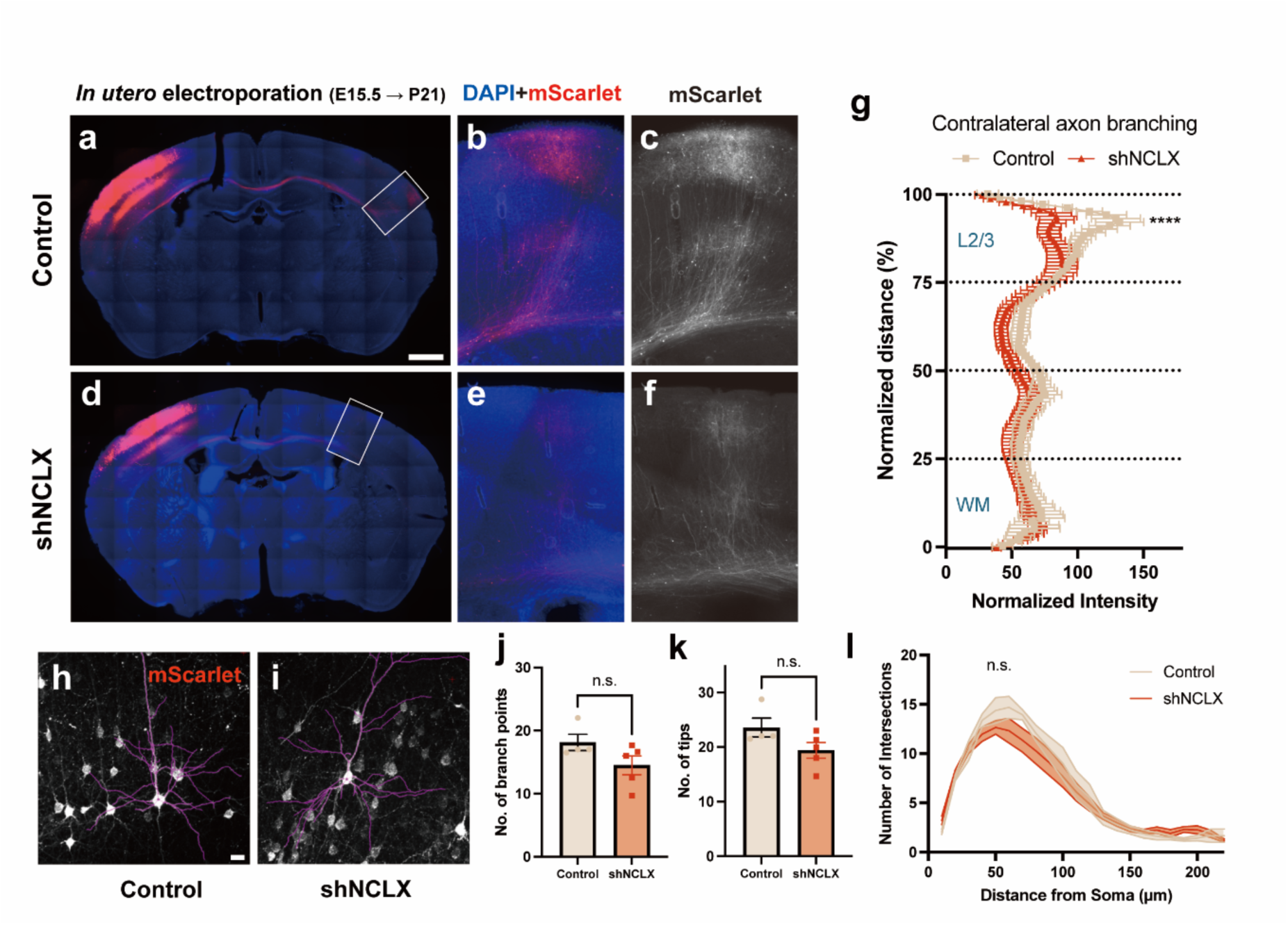
NCLX knockdown impairs axonal development, but does not significantly affect dendritic branching. (a, d) Representative image of a coronal section of P21 mouse brain following cortical IUE with mScarlet and either with control shRNA or shNCLX. (b, c, e, f) High magnification of the box in a, d. (g) Quantification of mScarlet fluorescence along the radial axis of the contralateral cortex reveals decreased contralateral terminal branching upon NCLX knockdown. (Two-way ANOVA, distance × intensity interaction F(100,299) = 1.604, p=0.001, Predicted LS mean fluorescence intensity at 93% normalized distance, Control 130.4, shNCLX 80.87, p<0.001). (h, i) Representative image of traced cortical neurons following IUE with mScarlet and either with control shRNA or shNCLX. (j-l) Quantification of dendritic branching using sholl analysis. Parameters analyzed include the number of branch points (j, Control 18.13±1.297, shNCLX 14.50±1.489, p=0.118) and number of tips (k, Control 23.56±1.733, shNCLX 19.40±1.439, p=0.104) the number of intersections (l, Two-way ANOVA, distance × intensity interaction: F(26, 77) = 1.409, p=0.126), were analyzed. n_Control_ = 4 mice, n_shNCLX_ = 5 mice. Data are presented as mean ± SEM. Two-way ANOVA with mixed-effects analysis with *post hoc* Sidak’s test was used for panel g and l, Unpaired t-test was used for panel j and k. ***p < 0.001, n.s., not significant. (a) Scale bar = 1mm, (h) Scale bar = 20μm

## Discussion

The structural contrast of neuronal mitochondria found in dendrites and axons and their functional relevance have been studied most often independently. Here, we demonstrate striking differences in the Ca^2+^ regulation by dendritic and axonal mitochondria in cortical pyramidal neurons. Axonal mitochondria maintain uptake even when ER-derived Ca^2+^ is depleted and display faster Ca^2+^ release, in contrast to dendritic mitochondria, which show strong ER dependence and slower Ca^2+^ release. These differences arise from distinct mitochondrial Ca^2+^ regulatory molecular compositions within each compartment, with axons showing higher levels of NCLX, MICU1 and MICU2. Importantly, this asymmetry is conserved in hippocampal pyramidal neurons, indicating that this compartmental difference is a generalizable feature of excitatory neurons. MICU1 or MICU2 overexpression enhanced Ca^2+^ uptake only in dendrites, while NCLX knockdown selectively slowed mitochondrial Ca^2+^ release in axons. Finally, NCLX deficiency significantly impaired axonal, but not dendritic branching, and MICU1 overexpression significantly increased dendritic mitochondrial vulnerability upon excitotoxicity, highlighting distinct physiological consequences of a compartment-specific requirement for mitochondrial Ca^2+^ regulation (Fig.7).

**Fig. 7.**
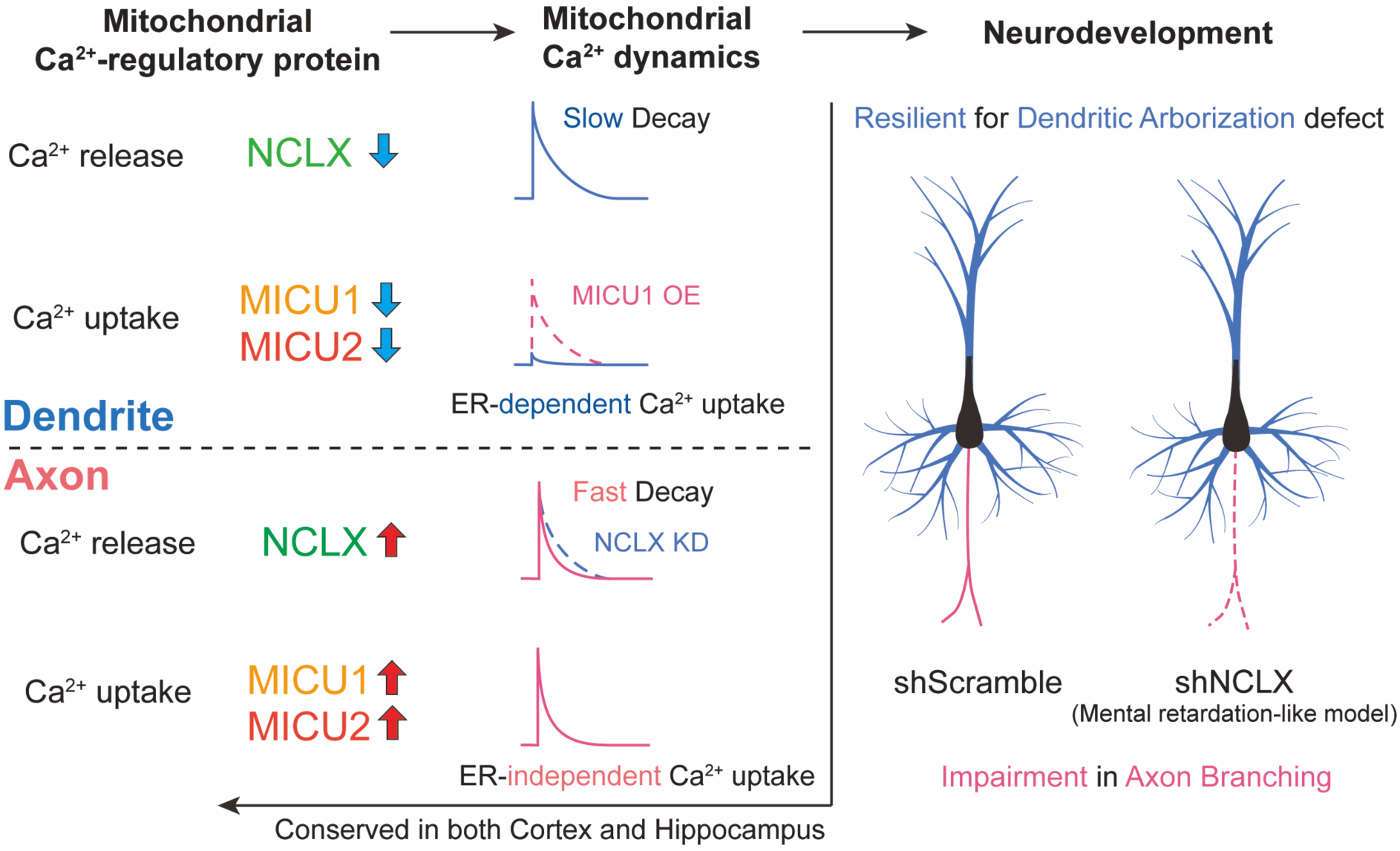
Compartment-specific asymmetry in mitochondrial Ca^2+^ regulation and functional properties. In dendrites (upper), synaptic input promotes ER-stored Ca^2+^ transfer to mitochondria. In contrast to axonal mitochondria, lower MICU1 and MICU2 level leads ER-dependent mitochondrial Ca^2+^ uptake, with slower mitochondrial Ca^2+^ efflux due to low NCLX expression. In axons (lower), synaptic input drives both ER and mitochondrial Ca^2+^ uptake. High NCLX expression in the axon facilitates rapid mitochondrial Ca^2+^ clearance, while axon-enriched MICU1 and MICU2 support ER-independent Ca^2+^ uptake. NCLX knockdown delay the rapid Ca^2+^ release in axonal mitochondria and leads axon branching impairment, while dendritic arborization was relatively resilient.

Previous study demonstrates that overexpression of mental retardation-associated NCLX mutant mimics the mitochondrial Ca^2+^ dynamics observed in NCLX KO hippocampal neurons, causing delayed Ca^2+^ release from the mitochondrial matrix ^36^. In addition, both presynaptic cytosolic Ca^2+^ concentration and neurotransmitter release are reduced in these KO neurons. Our NCLX knockdown axonal mitochondria display prolonged matrix Ca^2+^ decay, which may lead similar deficits in presynaptic release. Such altered presynaptic function could, in turn, impair axon branching development, a process known to be highly activity-dependent ^37^.

Loss-of-function mutations in *MICU1* are associated with abnormalities in brain development, learning difficulties, and movement disorders ^38–40^. Similarly, null mutation in *MICU2* has also been found in patients having neurodevelopmental disorders with severe cognitive impairment and spasticity ^41^. However, the progression of cellular defects by these mutations, particularly at the neuronal level, remains unclear. Defective mitochondrial Ca^2+^ uptake can elevate presynaptic Ca^2+^ and alter release properties, which can affect axon branching ^4, 42, 43^. Therefore, as observed in NCLX-deficient neurons, MICU1- or MICU2-mutatant neurons may exhibit more significant axonal than dendritic abnormalities.

Beyond neurodevelopment, mitochondrial Ca^2+^ dysregulation is also strongly related to neurodegenerative diseases ^11, 16, 17^. A former investigation showed that Parkinson’s disease-derived LRRK2 mutant increased mitochondrial Ca^2+^ uptake via MCU and MICU1 upregulation, resulting in dendritic shortening ^15^. In addition, restoring the mitochondrial Ca^2+^ homeostasis by overexpressing the NCLX active form rescued dendritic length, highlighting the importance of balanced mitochondrial Ca^2+^ dynamics for both axonal and dendritic arborization. Therefore, the compartment-specific enrichment of mitochondrial Ca^2+^ regulators may give asymmetric impacts on each domain.

MICU1 plays a dual role in regulating mitochondrial Ca^2+^ uptake, preventing excessive Ca^2+^ entry at low cytosolic Ca^2+^ concentrations while enhancing Ca^2+^ uptake at higher concentrations ^24, 44^. Interestingly, both silencing and overexpression of MICU1 leads to mitochondrial Ca^2+^ overloading, which triggers cell death ^25, 45, 46^. In our study, MICU1 overexpression enabled ER-independent Ca^2+^ uptake and led to rapid mitochondrial depolarization under excitotoxic conditions (Extended Data Fig. 7). Taken together, asymmetric distribution of MICU1 in neurons is critical not only for defining the source of Ca^2+^ entry but also protecting mitochondrial integrity and cell viability.

Based on previous studies, we propose additional functional implications of the distinct mitochondrial Ca^2+^ regulation observed in axonal and dendritic compartments:

(1) The differences in mitochondrial Ca^2+^ dynamics between axons and dendrites could be attributed to the different energy consumption requirements of each compartment. Synapses consume a substantial amount of ATP, with the dendrite utilizing a larger proportion compared to the axon ^47, 48^. This implies that ATP production might also be asymmetric in axons and dendrites. Given the critical role of mitochondrial Ca^2+^ in ATP production ^11^, our findings that distinct Ca^2+^ dynamics in axonal and dendritic mitochondria (Fig. 1) could be interpreted as these differences may be due to the unique energy needs of each compartment. Consequently, the slower decay of Ca^2+^ in dendritic mitochondria may enhance ATP generation to meet higher energy demands. ATP is also crucial for presynaptic function, with presynaptic mitochondria suggested to generate ATP in response to Ca^2+^ influx ^23^. However, glycolysis can also provide energy at presynaptic sites in multiple species ^49, 50^. This process could respond faster upon action potential arrival than oxidative phosphorylation and support mitochondria-free presynapses, which constitute more than half of excitatory presynaptic boutons. Intriguingly, a recent preprint study suggests that axonal mitochondria consume the ATP rather than generating it unlike dendritic mitochondria, possibly due to complex V hydrolysis ^51^.

(2) Compartment-specific mitochondrial Ca^2+^ dynamics may finely modulate synaptic communication. Mitochondrial Ca^2+^ buffering affects cytosolic Ca^2+^ concentration and this can contribute to Ca^2+^-activated K^+^ channel (K_Ca_) activity ^52^. Interestingly, K_Ca_ channels are expressed in a compartment-specific manner ^53^. The activity of large-conductance Ca^2+^-activated K^+^ channels (BK channels), which are enriched in axons, is critical for spike fidelity ^54^. A rapid Ca^2+^ release from axonal mitochondria may lead to a cytosolic Ca^2+^ transient that enhances BK channel activity, thus supporting spike fidelity in the axon. In contrast, some of small-conductance Ca^2+^-activated K^+^ channels like SK1 and SK2, are enriched in soma and dendrites ^53^. Given that dendritic mitochondria release Ca^2+^ at slower rates, SK channels may be activated more slowly. This could prevent excessive hyperpolarization of the membrane potential during synaptic events like burst firing, thereby maintaining sensitivity to subsequent synaptic events.

Further investigation is required to elucidate potential mechanisms underlying the polarized distribution of mitochondrial components. We speculate on multiple models, including early mitochondrial sorting in soma or distinct transport of mitochondrial protein or mRNA in a compartment-dependent manner. A previous report indicates that upon entry into the axon, mitochondria are already smaller compared to dendritic mitochondria ^1^, although the sorting mechanism remains unidentified. Then, NCLX and MICU1 may be enriched in the small mitochondria in the soma and recognized by mitochondrial motor adaptor proteins like TRAK1, which preferentially transport them to the axon ^55^. Alternatively, axon-targeted transport of NCLX and MICU mRNA or protein is also conceivable. Given that 99% of mitochondrial proteins are encoded in nuclear DNA, their conveyance is critical for the assembly of functional mitochondria. In addition, local translation of mRNA along the axon has long been studied and mitochondrial protein-encoding mRNAs have been detected in axons ^56–58^.

Overall, our study unveiled novel cellular mechanisms specifying neuronal polarized function via compartment-dependent mitochondrial Ca^2+^ regulation. These findings shine lights on the importance of mitochondrial molecular variability for physiological and pathophysiological studies.

## Data availability

All data reported in this paper will be shared by the corresponding authors upon request. This paper does not report original code. Any additional information required to reanalyze the data reported in this work paper is available from the corresponding authors upon request.

## Acknowledgements

We thank Tommy Lewis, Julien Courchet, Yusuke Hirabayashi, and Franck Polleux for discussions and critical feedback on the manuscript. We also thank all members of the Kwon lab and collaborators for feedback and discussion along the way. This research was supported by the National Research Foundation of Korea (NRF) grant funded by the Korean government (MSIT) (2020R1C1C1006386, 2022M3E5E8017395, RS-2023-00264980 to S-K. K.; RS-2021-NR061738 to D.C.J.) and KIST Program (2E33701 and 2E33721 to S-K. K.).

## Author contributions

Conceptualization: D.C.J., S.Y.K., W.S.K., S-K.K.; Methodology: D.C.J., S.Y.K., W.S.K., H.J., C.C., S.B., Y.C., S-K.K.; Investigation: D.C.J., S.Y.K., W.S.K., H.J., C.C., S.B.; Visualization: D.C.J., S.Y.K., W.S.K.; Funding acquisition: D.C.J., S-K.K.; Project administration: S-K.K.; Supervision: H.N.K., H.W.C., K.H., Y.C., S-K.K.; Writing – original draft: D.C.J., S.Y.K., W.S.K., S-K.K.; Writing – review & editing: D.C.J., S.Y.K., W.S.K., Y.C., S-K.K.

## Ethics declarations

Authors declare that they have no competing interests.

## Methods

### Ethical statement

All experiments were performed according to protocols approved by the Korea Institute of Science and Technology Institutional Animal Care and Use Committee (IACUC) (2023-047-2) and Institutional Biosafety Committee.

### Animals

Time-pregnant CD1 females were obtained from DBL. Littermates were randomly allocated to experimental groups without consideration of their sex at the moment of ex utero electroporation (E15.5).

### Constructs

pCAG::mito (COXVIII)-YFP, pCAG::mito (COXVIII)-mTagBFP2, pCAG::mTagBFP2 and pCAG:HA-mCherry were previously described.^1, 4^ pCAG::mScarlet was generated by replacing IRES-GFP from pCIG2 to mScarlet. For pCAG::mito-jGCaMP8m, we inserted jGCaMP8m from pGP-CMV-jGCaMP8m ^59^ (Addgene #162372) to pCAG::mito-mTagBFP2 by replacing mTagBFP2. pCAG::cyto-jGCaMP8s was generated by replacing IRES-GFP from pCIG2 to jGCaMP8s ^59^ (Addgene #162371). pCAG::vGlut1-jGCaMP8s was created by replacing GCaMP6s from pCAG::vGlut1-GCaMP6s ^60^ (Addgene #40753). The shNCLX-mTagBFP2 lentiviral shRNA constructs were created from pLKO.1-TRC cloning vector ^61^ (Addgene #10878). The target sequence used for shRNA knockdown of NCLX in mouse was as follows: 5’- CCAGACATCTTCAGTGCTTTA-3’. We replaced the Puromycin resistance sequence of scramble shRNA^62^ (Addgene #1864) with mTagBFP2 for the control vector used in shRNA experiments.

Control empty vector (pCAG::) was generated by deleting IRES-GFP sequence from pCIG2. pCAG::MICU1-mTagBFP2 and pCAG::MICU2-mTagBFP2 were created by inserting MICU1 (OriGene, MR207652) and MICU2 (OriGene, MR206892) into pCAG vector, respectively.

### Ex utero electroporation

A mixture of plasmid preparation (1-2 µg/µL) and 0.1% Fast Green (Sigma) was loaded in a glass capillary (WPI) and injected into the lateral ventricles of the isolated head of E15.5 mouse embryo. Then, the plasmid mixture was electroporated using an electroporator (ECM 830, BTX) with four pulses of 20 V for 100 ms with a 500 ms interval.

For cortical neuron targeting, the electroporation was carried out using gold-coated electrodes by placing the anode (positively charged electrode) on the side of DNA injection and the cathode (negatively charged electrode) on the bottom side of the head.

For hippocampal neuron targeting, the electroporation was performed using a triple-electrode setup. Two anodes were placed laterally on each the side of the head, and a single cathode was positioned rostrally at a 0° angle to the horizontal plane to target hippocampal progenitors.

### Primary neuron culture

Either embryonic mouse cerebral cortex or hippocampus was dissected in Hank’s buffered salt solution (HBSS, Invitrogen) supplemented with HEPES (10 mM, pH 7.4, Invitrogen), depending on the experimental requirements. Tissues were incubated in HBSS containing papain (Worthington) and DNaseⅠ (Sigma) and dissociated by pipetting. Dissociated cells (8.5×10^4^) were plated on poly-D-lysine (Sigma)/laminin (Gibco) coated 12 mm coverslip in Neurobasal media (Invitrogen) containing 2% B27, 1% Glutamax, 2.5% FBS and 1% penicillin/streptomycin (all supplements are from Invitrogen). After 5 to 7 days, the media was changed to an FBS-free supplemented Neurobasal medium. Neurons were maintained for 15–21 days in vitro (DIV) in a humidified 37 °C incubator with 5% CO_2_.

For Western blot experiments, dissociated cells (5.1×10^5^) were plated either on poly-D-lysine (1mg/ml, Sigma)-coated 24 mm Transwell® with 3.0 µm Pore Polycarbonate Membrane Insert (Corning) or PDL-coated 6-well plates in Neurobasal media (Invitrogen) containing 2% B27, 1% Glutamax, and 1% penicillin/streptomycin. Neurons were maintained for 10–11 days in vitro (DIV) in a humidified 37 °C incubator with 5% CO_2_.

### In utero electroporation

To label layer II/III cortical pyramidal neruons, in utero electroporation was done as previously described ^1^. A mixture of plasmids (1-2mg/ml) and 0.5% Fast green were injected to the lateral ventricle of E15.5 CD1 mouse embryos. Electroporations were performed with tweezer-type electrode. To label cortical progenitor cells, a positively charged part of electrode was placed on the top of DNA injection and a negatively charged part on the opposite side of the head. Four pulses of 40V for 50ms were used for electroporation. Mice were sacrificed 3 weeks after birth by transcardiac perfusion as previously described ^63^. Briefly, mice were anesthetized with 2.5 vol% isoflurane with 2ml/min oxygen and perfused transcardially with 4% PFA in PBS. Isolated brains were then post-fixed for 2 hours in 4% PFA and processed for further experiments. Brains were washed with PBS and sectioned using a vibratome (Leica VT1200) at either 130μm thickness. Sections were then washed three times with PBS and mounted on slides using VECTASHIELD® Vibrance™ Antifade Mounting Medium (Vector laboratories).

### HEK293FT cell transfection

Human Embryonic Kidney (HEK) 293FT cells (ThermoFisher) were maintained with Dulbeccos’s Modified Eagle Medium (Gibco) supplemented with 10% FBS (Gibco) and 1% Penicillin-Streptomycin (Gibco) in a humidified 37 °C incubator with 5% CO_2_. Cells were transfected via jetPrime (Polyplus) by following the manufacturer’s instructions. 3 days after transfection, cells were harvested for Western blot.

### Lentiviral production and infection

Lentiviruses were produced from HEK293FT cells by co-transfection with shuttle vectors, LP1, LP2, and VSV-G. jetPrime (Polyplus) was used as manufacturer’s protocol and 24 hours after transfection, media were changed with Neurobasal media, and 48 hours later, supernatants were harvested and centrifuged to remove cellular debris. 100-200μl of viral supernatants were added to primary cortical neurons at 3-5 DIV and harvested 10 days after.

### Western blotting

Proteins from whole cell fraction and axonal fraction were separately prepared from 3 µm porous membrane. To obtain the whole cell fraction, cold DPBS was applied on the upper side of the membrane and cells were harvested using cell scrapers (SPL). After a brief centrifuge, the supernatant was removed, and cells were lysed using RIPA buffer (50 mM Tris-HCl pH7.4, 150 mM NaCl, 1% NP40, 0.5% Sodium deoxycholate, 0.1% SDS) supplemented with the cocktail of protease inhibitor and phosphatase inhibitor at 4°C for 1h. Before collecting the axonal fraction, the upper side of the membrane was wiped thoroughly using cotton swap to prevent contamination of the whole cell fraction. The membranes were submerged in the same lysis buffer for the axonal fraction and vortexed for 5 min. After vortexing, membranes were discarded, and cell lysate was incubated at 4°C for 1 hour.

5 µg of total protein per well was loaded on 10-12% SDS-PAGE gels and transferred to nitrocellulose membrane (Merck). After transfer, membranes were blocked for 30 min with 5% skim milk (LPS Solution), followed by incubation with primary antibodies overnight at 4°C. The membranes were washed three times with 0.1% Tween20/TBS (TBST) and incubated with HRP-conjugated secondary antibodies (Abbkine) for 1 h at room temperature, followed by three washes with TBST. The bands were detected using ECL Western blotting substrate (Amersham) and then scanned by iBright CL750 Imaging System (Thermofisher).

The following primary antibodies were used for Western blotting in the study: MCU (Invitrogen, MA5-24702, 1:1000), MICU1 (Atlas antibodies, HPA037480, 1:1000), MICU2 (Abcam, ab101465, 1:2000), MICU3 (Invitrogen, PA5-55177, 1:2000), VDAC1 (Abcam, ab14734, 1:1000), NCLX (Abcam, ab83551, 1:1000), TOM40 (Proteintech, 18409-1-AP, 1:2000), Synaptophysin (Synaptic Systems, 101011, 1:5000), Histone H3 (Cell Signaling, 96C10, 1:1000), GluR2 (Sigma, MAB397, 1:1000), beta-actin (Abbkine, 1C7, 1:10000). All secondary antibodies were HRP-conjugated (Abbkine, A21020 and A21010) and used at a 1:10000 dilution.

### Immunocytochemistry

Cells were fixed for 12 min at room temperature in 4% paraformaldehyde (PFA, T&I) with 4% sucrose (Sigma) in PBS (Sigma) and then washed 3 times with PBS. Samples were permeabilized with 0.2% Triton X-100 (Sigma) in PBS and incubated for 30 min with 2.5% goat serum and 0.1% BSA (Sigma) in PBS to block nonspecific signals. Primary and secondary antibodies were diluted in the blocking buffer as described above. Primary antibodies were incubated overnight at 4°C, and secondary antibodies were incubated for 1 hour at room temperature. Coverslips were mounted on slides with VECTASHIELD® Vibrance™ Antifade Mounting Medium (Vector Laboratories).

The following primary antibodies were used for immunocytochemistry in the study: MCU (Novus, NBP2-76948, 1:200), HA (Biolegend, 901513, 1:500). All secondary antibodies were Alexa-conjugated (Invitrogen, A11003 and A21244) and used at a 1:1000 dilution.

### Analysis of axon branching and dendritic branching

For analysis of axon branching, brain sections at similar rostrocaudal level were selected, and axons on the contralateral side were evaluated. Quantification was performed by measuring relative fluorescence intensity along the radial axis of the contralateral cortex. To account for variability in transfection efficiency, the fluorescence intensity of contralateral axons was normalized to that of intensity of collosal axons within the same section ^1, 42^.

For analysis of dendritic branching, brain sections from the same mice used for axon branching analysis were examined. Transfected neurons in the ipsilateral cortex were analyzed using Simple Neurite Tracer (SNT) ^64^ plugin in ImageJ. Individual dendrites were manually traced, and the number of branching points and dendritic tips was measured. Sholl analysis was performed on the same traced neurons using concentric circles spaced at 10μm intervals to evaluate dendritic complexity relative to the distance of the soma.

### Imaging

Fixed samples were imaged on a Nikon A1R confocal microscope (60x objective NA 1.25). All equipment and solid state lasers (Coherent, 405, 488, 561, and 647 nm) were controlled via Nikon Elements software. Samples were visualized by Z-stacking that was later processed into Maximum Intensity Projection (MIP). MIP 2D images were then analyzed with Fiji (Image J).

For live imaging, electroporated cortical neurons were imaged at 15-21DIV with EMCCD camera (Andor, iXon Life 897) on Nikon microscope FN1 (60x objective NA1.0). 395, 470 and 555 nm Spectra X LED lights (Lumencor) were used for the light source. We used modified normal Tyrode’s solution as a bath solution containing the following (in mM): 145 NaCl, 2.5 KCl, 10 HEPES, 2 CaCl_2_, 1 MgCl_2_, 10 glucose, pH 7.4 at RT. The electrical stimulation is triggered by 1 ms current injections with a glass capillary electrode placed near the transfected axon, dendrite or soma. We applied 10 pulses (10Hz) with 30 µA using a stimulator (Model 2100, A-M systems) and imaged with 500 ms interval (2Hz) during 180 s for mito-jGCaMP8m and cyto-jGcaMP8s signals. To avoid the spontaneous activity of cultured neurons, we treated a small amount of tetrodotoxin (TTX, less than 50 nM). For blocking sarco/endoplasmic reticulum Ca^2+^-ATPase (SERCA) pump, we treated cyclopiazonic acid (CPA, 30 μM, Tocris) for longer than 5 min.

For Tetramethylrhodamine (TMRM, Invitrogen) imaging, cells were incubated with 50 nM TMRM for 10 min at 37 °C before imaging started. To investigate changes in mitochondrial membrane potential under excitotoxic conditions, 20 μM N-Methyl-D-aspartic acid (NMDA, Sigma) was added to the bath solution. Before NMDA treatment, cells were imaged with 500 ms interval for 5 min to assess the fluorescence baseline, then imaged for an additional 10 min after treatment.

### Statistics

Ca^2+^ imaging data was analyzed via Nikon Element and custom-built Python code. Western blot data was analyzed with iBright Analysis Software (Thermo Scientific). All statistical analysis was performed using the latest version of GraphPad Prism and Microsoft Excel. One- and Two-way ANOVA with post hoc tests were used for group comparisons. All graphs are shown as mean ± SEM, and asterisks *, ** and *** indicate p<0.05, p<0.01 and p<0.001, respectively. Detailed statistical methods and n for each experiment are written in the figure legends.

## Extended Data Figures

**Extended Data Fig. 1.**
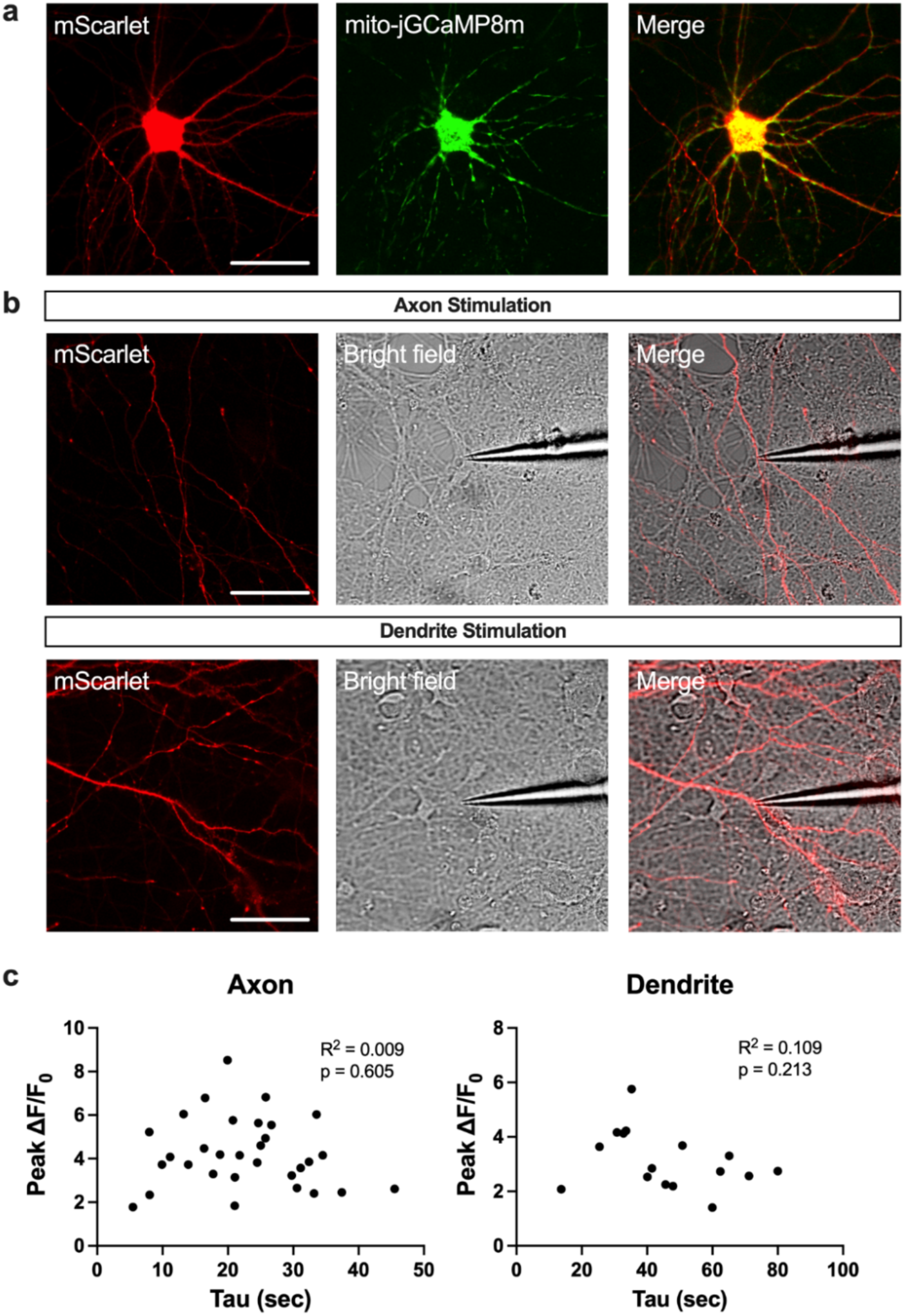
Representative images of recorded neurons and correlation between peak intensity and decay tau in mitochondria. (a) Representative images of recorded neurons. pCAG::mScarlet and pCAG::mito-jGCaMP8m were electroporated into the cortical pyramidal neurons and monitored at 17-21 DIV. Scale bar, 50 µm. (b) Representative images of axon (upper) and dendrite (lower) stimulation. The stimulation electrode (glass capillary) was placed <10 µm toward the z-axis from the target (middle, bright field). (c) There was no significant correlation between peak intensity and decay tau in both axonal (left) and dendritic (right) mitochondria (Axon p=0.605, Pearson r =-0.0967, R^2^=0.00935; Dendrite p=0.213, Pearson r =-0.330, R^2^=0.109).

**Extended Data Fig. 2.**
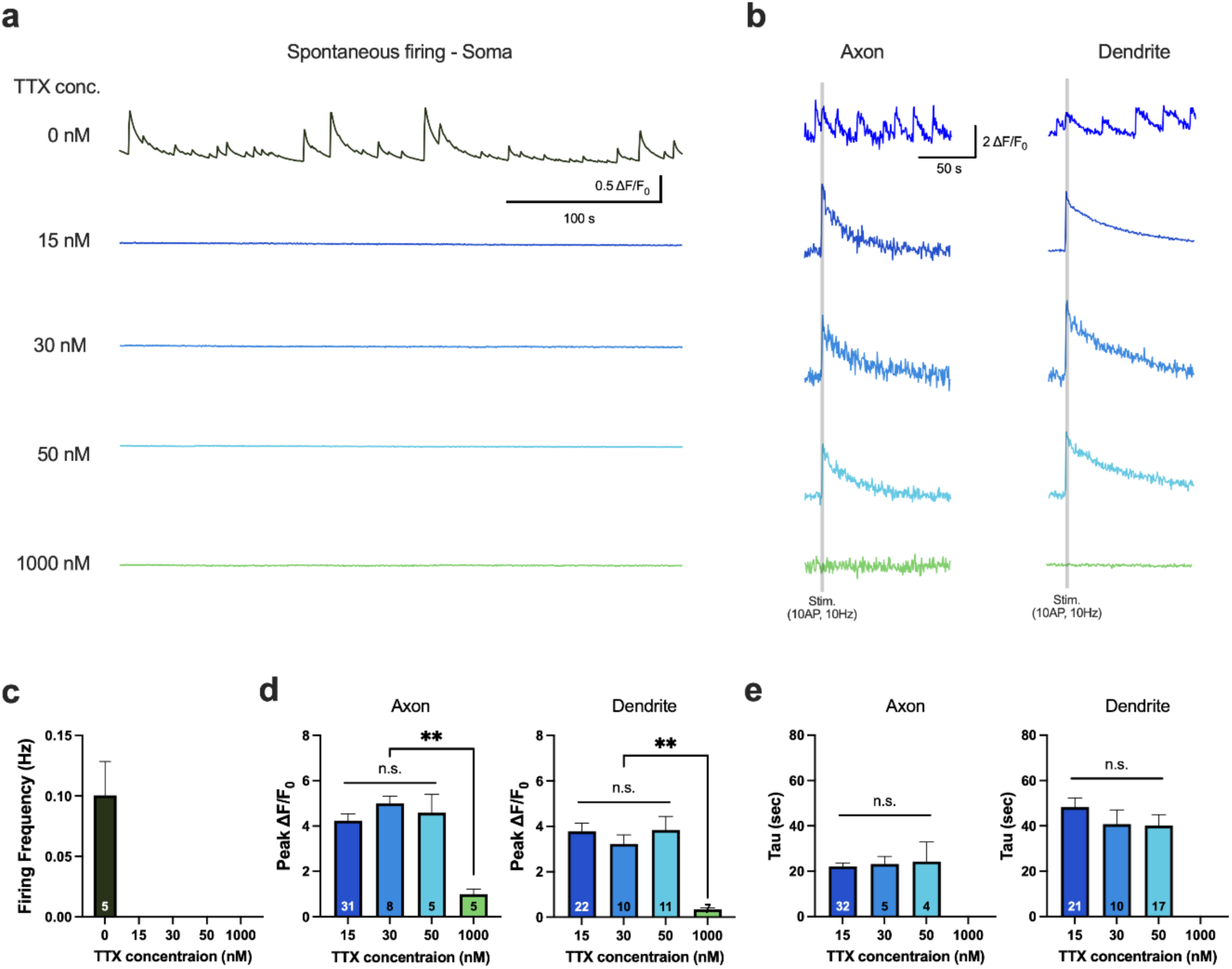
Mitochondrial Ca^2+^ transients under various tetrodotoxin (TTX) concentrations. (a, c) Spontaneous firing was observed with no TTX treatment (0 nM), but 15 nM or higher concentrations of TTX blocked the spontaneous firing. (b, d) Evoked Ca^2+^ responses in axonal and dendritic mitochondria were similar between 15 nM and 50 nM TTX (Axon, One-way ANOVA, F=8.69, p<0.001; 15nM 4.24±0.288, 30nM 4.99±0.319, 50nM 4.58±0.817, 1000nM 0.993±0.227; 15nM vs 30nM p=0.571, 15nM vs 50nM p=0.962, 15nM vs 1000nM p<0.001, 30nM vs 50nM p=0.961, 30nM vs 1000nM p<0.001, 50nM vs 1000nM p=0.002; Dendrite, One-way ANOVA, F=9.31, p<0.001; 15nM 3.78±0.364, 30nM 3.23±0.394, 50nM 3.84±0.592, 1000nM 0.339±0.0639; 15nM vs 30nM p=0.798, 15nM vs 50nM p>0.999, 15nM vs 1000nM p<0.001, 30nM vs 50nM p=0.808, 30nM vs 1000nM p=0.003, 50nM vs 1000nM p<0.001). The Ca^2+^ response was relatively smaller without TTX treatment, and almost no response was observed at 1000 nM TTX. (e) TTX treatment did not affect mitochondrial Ca^2+^ decay tau (Axon, One-way ANOVA, F=2.06, p=0.122; 15nM 22.2±1.44, 30nM 23.2±3.38, 50nM 24.3±8.56, 1000nM 0.00±0.00; 15nM vs 30nM p=0.996, 15nM vs 50nM p=0.971, 15nM vs 1000nM p=0.095, 30nM vs 50nM p=0.998, 30nM vs 1000nM p=0.111, 50nM vs 1000nM p=0.097; Dendrite, One-way ANOVA, F=2.32, p=0.088; 15nM 48.3±4.07, 30nM 40.7±6.26, 50nM 40.1±4.88, 1000nM 0.00±0.00; 15nM vs 30nM p=0.743, 15nM vs 50nM p=0.577, 15nM vs 1000nM p=0.086, 30nM vs 50nM p>0.999, 30nM vs 1000nM p=0.204, 50nM vs 1000nM p=0.200). One-way ANOVA with *post hoc* Tukey’s test was used for panels d and e. Mitochondrial numbers (1 mitochondrion image is analyzed from each axon or dendrite) are annotated at the bottom of bar graphs. Somal, Axonal and dendritic mitochondrial Ca^2+^ dynamics of cortical pyramidal neurons were monitored using mito-jGCaMP8m at 15-21 DIV. Data are presented as mean ± SEM. **p < 0.01. ***p < 0.001. n.s., not significant.

**Extended Data Fig. 3.**
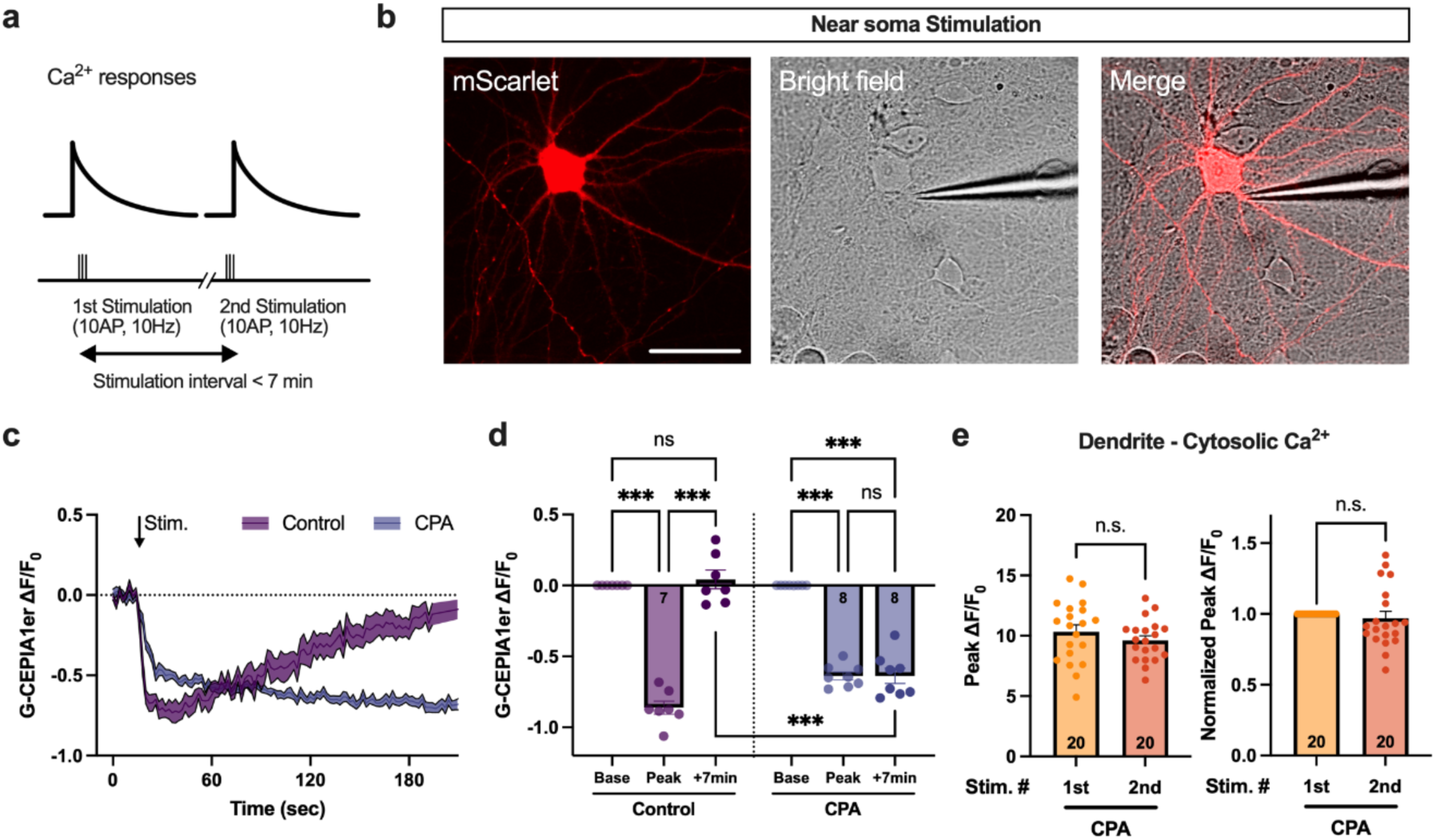
Stimulation protocol and representative images of recorded neurons. Axonal and dendritic mitochondrial Ca^2+^ dynamics of CPA treated cortical pyramidal neurons were monitored using mito-jGCaMP8m at 15-21 DIV. (a) Schematic diagram of the stimulation protocol. Each stimulation consists of 10 pulses at a 10 Hz frequency. After the first stimulation, there is a ∼7- min interval before the second stimulation. (b) A representative image of the stimulation method. The stimulation electrode (glass capillary) was placed <10 µm toward the z-axis from the target (middle, bright field). (c,d) Dendritic ER Ca^2+^ dynamics was monitored by G-CEPIA1er with CPA treatment. (c) The control group showed a rapid signal reduction after stimulation, but the reduced signal almost recovered around 3 min after stimulation (n=7). In contrast, the CPA group showed a continuous reduction of the signal after stimulation (n=8). (d) The signal from the control group was significantly reduced at peak and fully recovered within 7-min after stimulation. In contrast, the reduced signal remained constant until 7-min after stimulation in the CPA-treated group. At 7-min after stimulation, the fluorescence signal between the control and CPA group was significantly different (One-way ANOVA, F=99.9, p<0.001; Control Base −0.0001±0.00002, Control Peak −0.861±0.0462, Control+7min 0.0419±0.0658; Control Base vs Control Peak p<0.001, Control Peak vs Control+7min p<0.001, Control Base vs Control+7min p=0.990; CPA Base −0.0003±0.00002, CPA Peak −0.638±0.0276, CPA+7min 0.637±0.0533; CPA Base vs CPA Peak p<0.001, CPA Peak vs CPA+7min p>0.999, CPA Base vs CPA+7min p<0.001, Control+7min vs CPA+7min p<0.001). (e) Cytosolic Ca^2+^ levels from the dendrite were not different between the first and the second stimulation under CPA treated condition (Peak, 1^st^ stim. 10.3±0.576, 2^nd^ stim. 9.61±0.384, p=0.303; Normalized peak, 1^st^ stim. 1.000±0.000, 2^nd^ stim. 0.969±0.0488, p=0.531). One-way ANOVA with *post hoc* Sidak’s test was used for panel D. Unpaired t-test was used for panel e. Dendritic numbers are indicated at the bottom of the bar graph. Data are presented as mean ± SEM. n.s., not significant. Scale bar, 50 µm.

**Extended Data Fig. 4.**
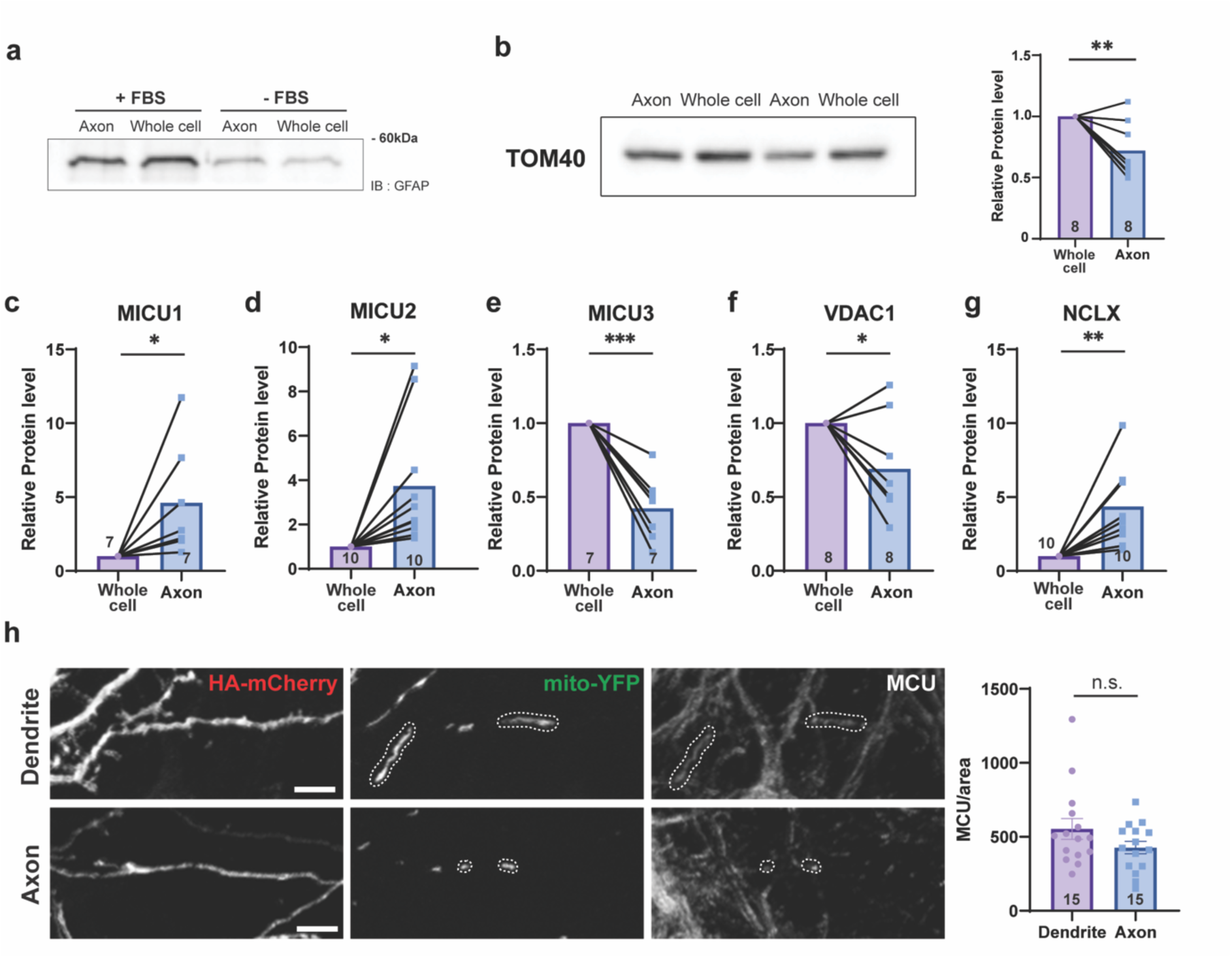
Relative expression levels of mitochondrial Ca^2+^ regulatory proteins. (a) Culture media without FBS shows lower level of GFAP (astrocyte marker) in both axonal and whole cell fraction at 10 DIV. (b) Western blot result of TOM40 (Axon 0.721±0.081, p=0.011). (c-g) Western blot results of mitochondrial Ca^2+^ regulatory protein levels without MCU normalization. MICU1 (c, Axon 4.622±1.441, p=0.046), MICU2 (d, Axon 3.733±0.901, p=0.014) and NCLX (g, Axon 4.369±0.830, p=0.003) levels are higher in axonal fraction. In contrast, MICU3 (e, Axon 0.423±0.084, p<0.001) and VDAC1 (f, Axon 0.691±0.119, p=0.036) showed lower levels. (h) Endogenous MCU levels in dendritic and axonal mitochondria of cortical pyramidal neurons were quantified using immunostaining at 10 DIV. The MCU expression in dendritic and axonal mitochondria is similar. (Dendrite 427.5±42.44, Axon 552.9±69.86, p=0.136). Relative protein levels were normalized to values from the whole cell fraction. Paired t-test was used for panels b to g. Unpaired t-test was used for panel h. Sample numbers are annotated at the bottom of bar graphs. Data are presented as mean ± SEM. *p < 0.05. **p < 0.01. ***p < 0.001. n.s., not significant. Scale bar, 10 µm

**Extended Data Fig. 5.**
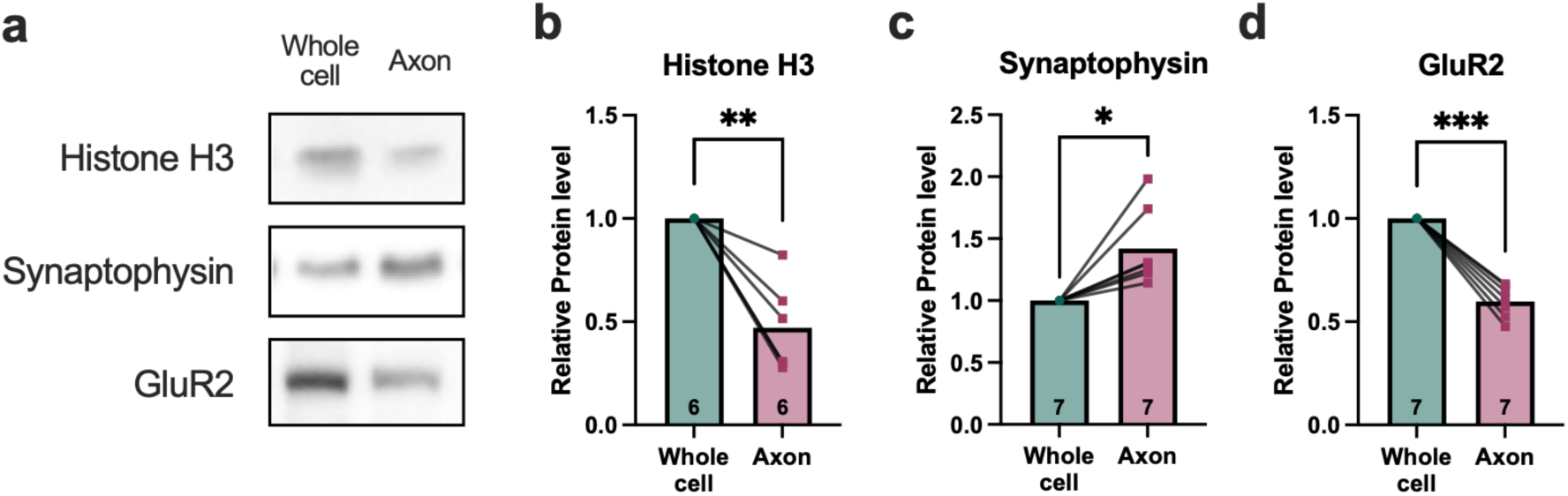
Validation of axon isolation in primary hippocampal culture. Separation of whole cell and axonal fraction is confirmed by synaptophysin (presynapse/axon), histone H3 (nuclear/cell body), and GluR2 (postsynapse/dendrite) levels. (a) Representative Western blot images. Bar graph showing synaptophysin (c, Axon 1.42±0.119, p=0.012) level is significantly higher in the axonal fraction, but not histone H3 (b, Axon 0.471±0.0887, p=0.002)) and GluR2 (d, Axon 0.599±0.0296, p<0.001)). Relative protein levels were normalized to values from the whole cell fraction. Paired t-test. Sample numbers are annotated at the bottom of bar graphs. Data are presented as mean ± SEM. *p < 0.05. **p < 0.01. ***p < 0.001.

**Extended Data Fig. 6.**
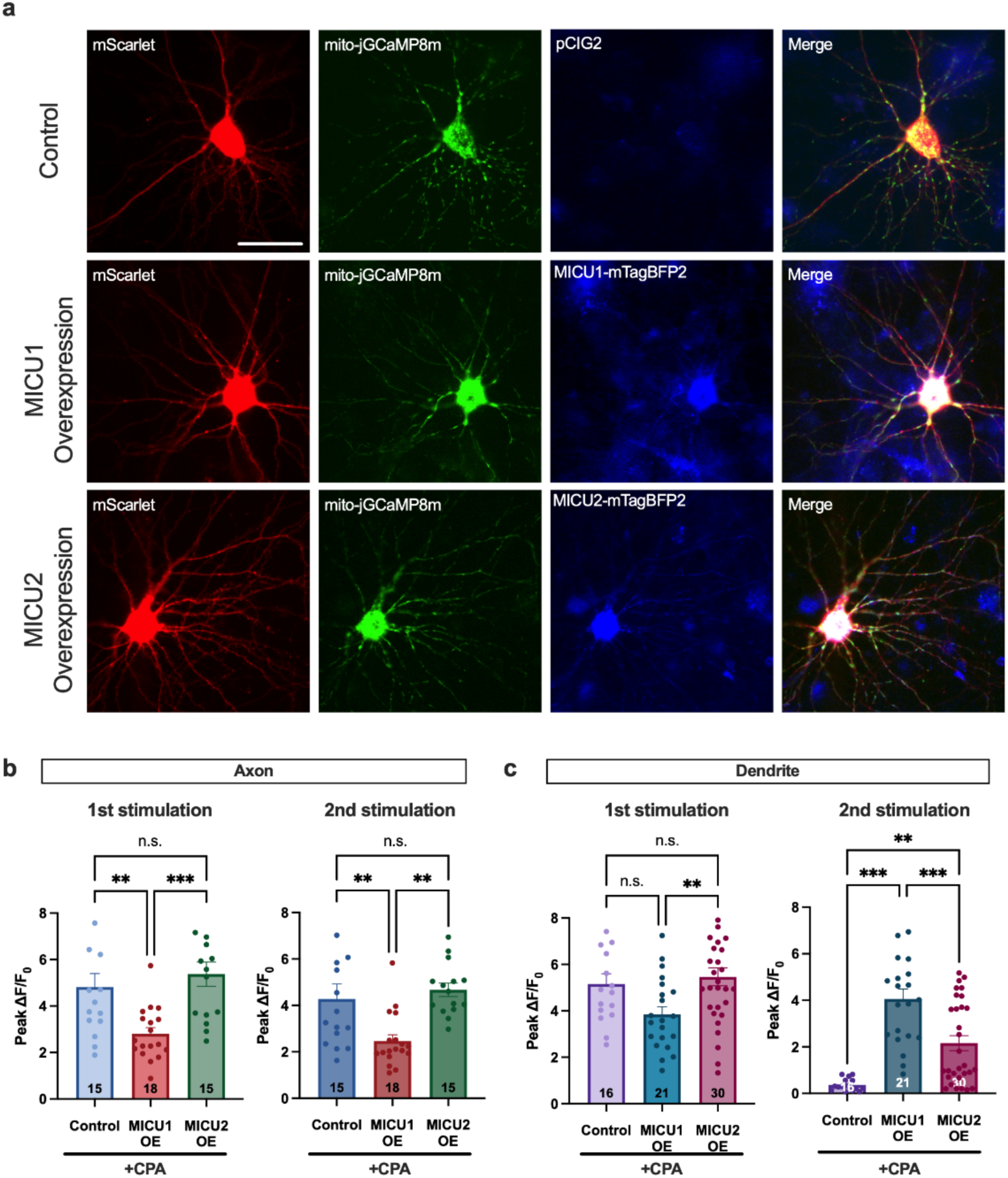
Validation of MICU1 and MICU2 overexpression and representative images of recorded neurons. (a) Representative images of recorded neurons. pCAG::mScarlet, pCAG::mito-jGCaMP8m and pCIG2 or pCAG::MICU1-mTagBFP2 or pCAG::MICU2-mTagBFP2 were electroporated into the cortical pyramidal neuron. (b, c) The peak intensity from axonal and dendritic mitochondria. (b) In axonal mitochondria, peak intensity was reduced and increased in the MICU1 and MICU2 overexpression groups, respectively. In the MICU1 overexpression group, the significant reduction was shown at the 2^nd^ stimulation, but in the MICU2 overexpression group, there was a tendency of increased Ca^2+^ uptake at the 1^st^ stimulation (1^st^ stimulation, One-way ANOVA, F=9.21, p<0.001; Control 4.82±0.581, MICU1 OE 2.80±0.266, MICU2 OE 5.37±0.525; Control vs MICU1 OE p=0.008, Control vs MICU2 OE p=0.686, MICU1 OE vs MICU2 OE p<0.001; 2^nd^ stimulation, One-way ANOVA, F=8.24, p<0.001; Control 4.28±0.643, MICU1 OE 2.46±0.275, MICU2 OE 4.67±0.289; Control vs MICU1 OE p=0.010, Control vs MICU2 OE p=0.799, MICU1 OE vs MICU2 OE p=0.001) (c) In dendritic mitochondria, both MICU1 and MICU2 overexpression enhanced Ca^2+^ uptake at the 2^nd^ stimulation, but in different amounts. Compared to the control group, the MICU1 overexpression group showed considerably higher peak intensity after ER-Ca^2+^ depletion, but partial enhancement by MICU2 overexpression was observed (1^st^ stimulation, One-way ANOVA, F=4.85, p=0.011; Control 5.16±0.441, MICU1 OE 3.85±0.338, MICU2 OE 5.47±0.385; Control vs MICU1 OE p=0.096, Control vs MICU2 OE p=0.854, MICU1 OE vs MICU2 OE p=0.009; 2^nd^ stimulation, One-way ANOVA, F=23.6, p<0.001; Control 0.364±0.0639, MICU1 OE 4.06±0.431, MICU2 OE 2.16±0.324; Control vs MICU1 OE p<0.001, Control vs MICU2 OE p=0.002, MICU1 OE vs MICU2 OE p<0.001). One-way ANOVA with post hoc Tukey’s test was used for panels c, d. Sample numbers are annotated at the bottom of bar graphs. Scale bar, 50 µm.

**Extended Data Fig. 7.**
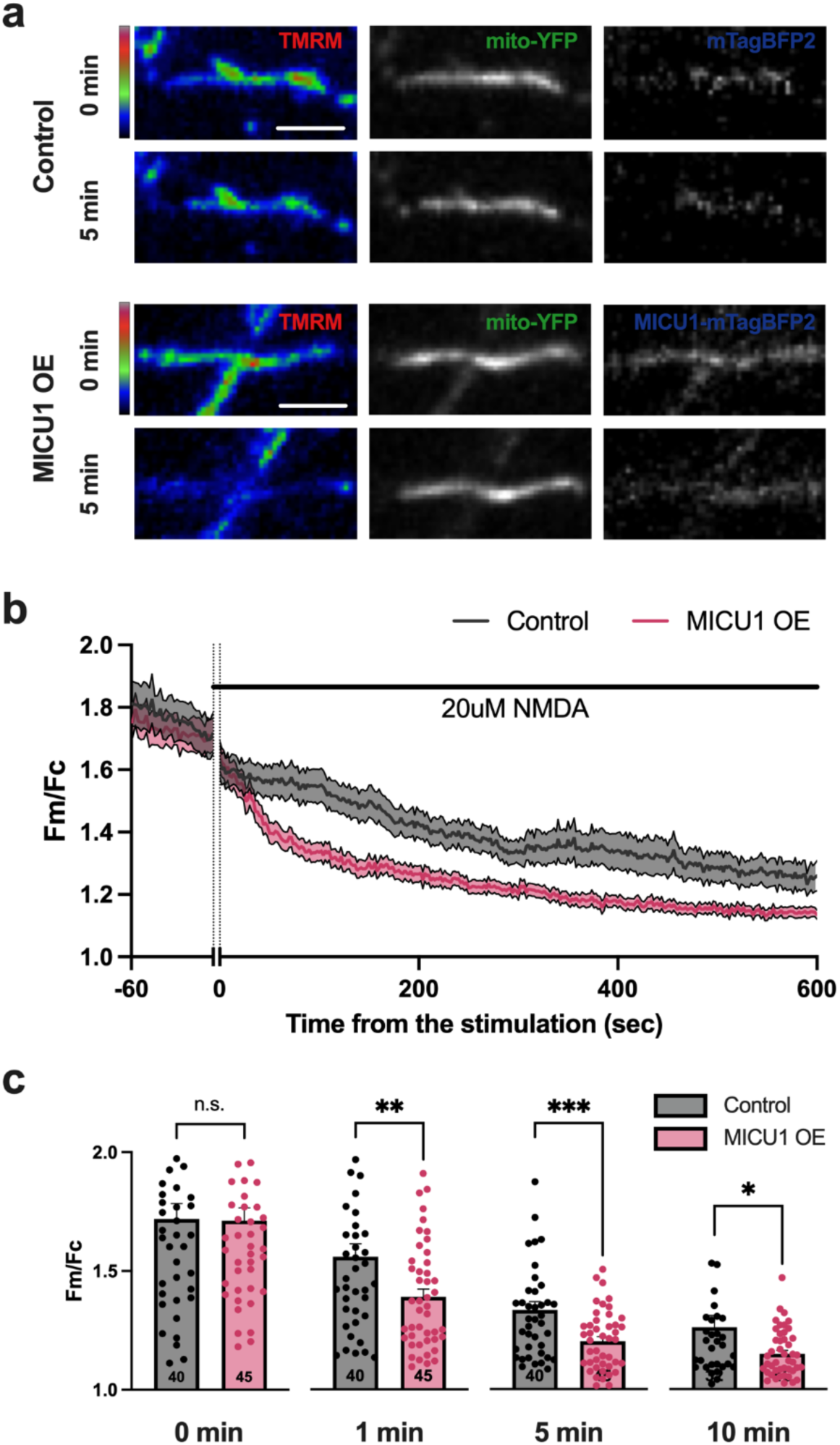
MICU1 overexpression accelerates the decrease in mitochondrial membrane potential under excitotoxic conditions. (a) Representative images from control and MICU1 overexpression groups. (b) Time course of mitochondrial membrane potential, represented as F_mitochondria_/F_cytoplasm_(Fm/Fc), in response to 20 µM NMDA stimulation in control and MICU1 overexpression (MICU1 OE) conditions. (c) Quantification of Fm/Fc at 0, 1, 5, and 10 minutes post-stimulation, showing significantly lower values in MICU1 OE compared to control at 1, 5, and 10 minutes. No significant difference was observed at 0 minutes. (40 mitochondria from 40 dendrites for control, 45 mitochondria from 45 dendrites for MICU1 OE; 0 min, Control 1.72±0.0652, MICU1 OE 1.71±0.0529, p=0.940; 1 min, Control 1.56±0.0540, MICU1 OE 1.39±0.0328, p=0.008; 5 min, Control 1.33±0.0346, MICU1 OE 1.20±0.0191, p<0.001; 10 min, Control 1.26±0.0487, MICU1 OE 1.15±0.0150, p=0.013). Data are presented as mean ± SEM. *p < 0.05, **p < 0.01, ***p < 0.001, n.s., not significant. Scale bar 5µm.

**Extended Data Fig. 8.**
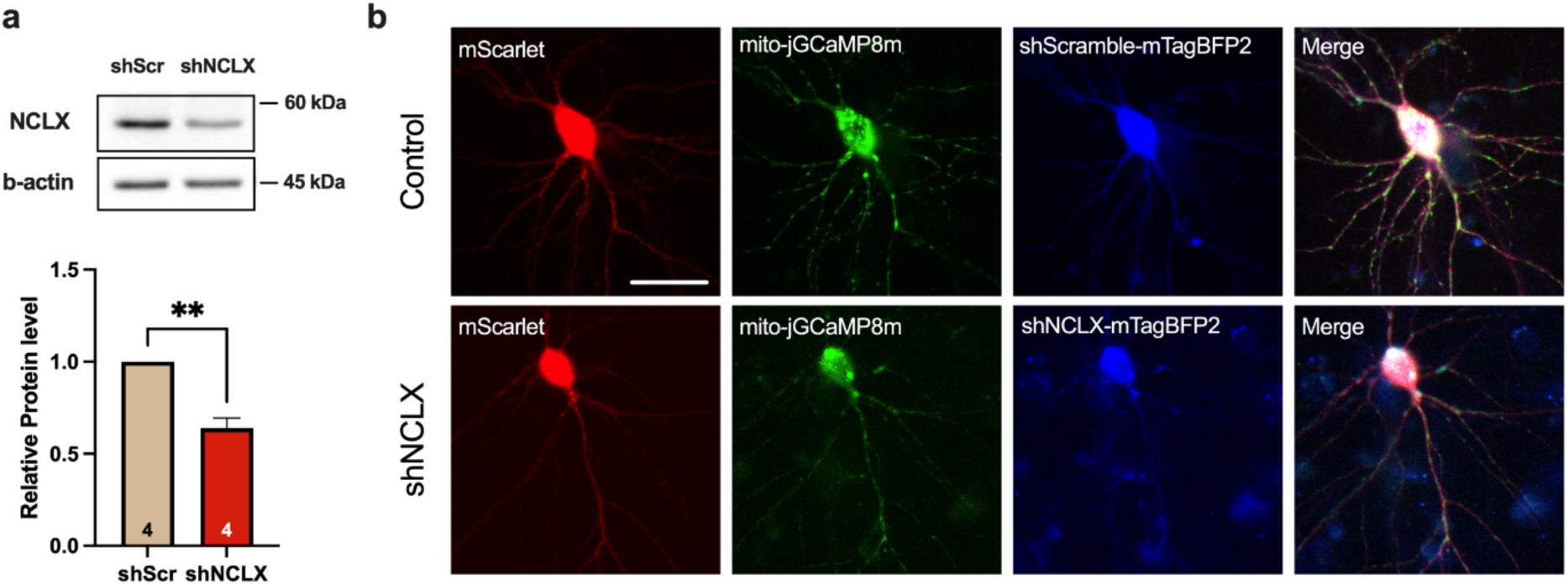
NCLX knockdown validation and representative images of recorded neurons. (a) Representative Western blot image from shScramble and shNCLX groups. Cortical neurons were infected by shScramble or shNCLX lentiviruses at 3-5 DIV and incubated for 10 days. Endogenous NCLX protein level is significantly reduced by NCLX knockdown. Relative protein level was normalized to values from the shScramble. (shScarmble 1.000±0.000, shNCLX 0.640±0.543, p=0.007) (b) Representative images of recorded neurons. pCAG::mScarlet, pCAG::mito-jGCaMP8m, and pLKO-shScramble-mTagBFP2 or pLKO.1-shNCLX-mTagBFP2 were electroporated into the cortical pyramidal neuron. Paired t-test was used for bar graph in panel a. Sample numbers are annotated at the bottom of bar graph. Data are presented as mean ± SEM. **p < 0.01. Scale bar, 50 µm

